# Exploring the effect of mechanical anisotropy of protein structures in the unfoldase mechanism of AAA+ molecular machines

**DOI:** 10.1101/2022.04.06.487390

**Authors:** Rohith Anand Varikoti, Hewafonsekage Yasan Y. Fonseka, Maria S. Kelly, Alex Javidi, Mangesh Damre, Sarah Mullen, Jimmie L. Nugent, Christopher M. Gonzales, George Stan, Ruxandra I. Dima

## Abstract

Essential cellular processes of microtubule disassembly and protein degradation, which span lengths from tens of *μ*m to nm, are mediated by specialized molecular machines with similar hexameric structure and function. Our molecular simulations at atomistic and coarse-grained scales show that both the microtubule severing protein spastin and the caseinolytic protease ClpY, accomplish spectacular unfolding of their diverse substrates, a microtubule lattice and dihydrofolate reductase (DHFR), by taking advantage of mechanical anisotropy in these proteins. By considering wild-type and variants of DHFR, we found that optimal ClpY-mediated action probes favorable orientations of the substrate relative to the machine. Unfolding of wild-type DHFR involves strong mechanical interfaces near each terminal and occurs along branched pathways, whereas unfolding of DHFR variants involves softer mechanical interfaces and occurs through single pathways, but translocation hindrance can arise from internal mechanical resistance. For spastin, optimum severing action initiated by pulling on a tubulin subunit is achieved through the orientation of the machine versus the substrate (microtubule lattice). Moreover, changes in the strength of the interactions between spastin and a microtubule filament, which can be driven by the tubulin code, lead to drastically different outcomes for the integrity of the hexameric structure of the machine.

## 1. Introduction

The AAA+ (ATPases associated with diverse cellular activities) protein superfamily comprises large biomolecular machines that perform mechanical action during fundamental processes in the life of a cell such as protein degradation (caseinolytic proteases ClpA, ClpX or ClpY), disassembly of toxic protein aggregates (ClpB or heat-shock protein Hsp104), DNA replication, severing of microtubules (MTs) during mitosis (katanin and spastin), cilia and flagella motions, and neuronal transmission [1–4]. Three-dimensional structures of these molecular machines reveal single– (ClpX, ClpY, katanin, spastin) or double–ring (ClpA, ClpB, Hsp104) hexameric assemblies that enclose a narrow central channel with a diameter of ~ 2 nm [5–8]. Their activity is provided by one or two conserved nucleotide– binding domains (NBDs), or AAA domains, that undergo large–scale conformational transitions driven by ATP hydrolysis in the individual monomers [9,10]. AAA rings have distinct sequence and possess three-dimensional structural characteristics that underlie their classification into clades, such as clade 3 for microtubule-severing proteins, clade 5 for ClpX or ClpY, and clades 3 and 5, respectively, for the two rings of ClpA or ClpB. Mechanical force is applied by AAA+ machines within the narrow central channel, as the substrate protein (SP) is ensnared by a set of protruding loops that execute ~ 1 nm axial motions [11–15]. The non-planar aspect of the hexameric assembly suggested a “hand-over-hand” SP translocation mechanism driven by nonconcerted ATP hydrolysis in each ring, comprising SP grip and release by successive central channel loops of each AAA domain [15,16]. On faster timescales (*μ*s - ms), thermal motion of the loops is proposed to enable rapid translocation of polypeptide chains through a Brownian ratchet mechanism [17,18].

Microtubules (MTs), which are polymeric assemblies of tubulin dimers, are the longest and stiffest filaments in the cell, with persistence lengths of mm [19], Young’s moduli of ~ 1 GPa (axial), ~ 10 MPa (circumferential), and a shear modulus of ~ 1 GPa [20,21]. During numerous cellular processes, such as mitosis and meiosis, the cell needs to cut down these long filaments at locations distant from their ends using microtubules severing enzymes, which function as hexamers. In severing proteins the AAA domain is connected via a long and flexible linker to an N-terminal domain that is known to bind microtubules with low affinity [22,23]. The N-terminal domain contains a Microtubule Interacting and Trafficking (MIT) domain, which consists of a three-helix bundle (PDB code 2RPA) [24]. The six arms, which contain the flexible linkers and the MIT domains, and are directed outward from the AAA motor region, are likely to be used by the enzyme to dock onto the MT [25,26]. Importantly, experiments found that just the AAA domain has no measurable severing activity, especially in katanin [25,27]. Thus, for both katanin and spastin the AAA domain and the linker are required for severing [25,27,28]. The orientation of the machine on a MT filament is one of the outstanding unknowns in the field. Namely, major barriers in finding the mechanism of substrate engagement by severing enzymes [26] are the lack of structures available for complexes between severing enzymes and MTs, and the lack of understanding how the MIT domain and the flexible linker recognize the MT [29,30].

Often, the rate-limiting step of the remodeling action of AAA+ machines corresponds to the process of SP unfolding, which is modulated by the protein structure, either through the topology near the terminal engaged by the machine or through internal features. In stringent cases, protein degradation may stall altogether due to strong mechanical resistance associated with complex structure, such as in knotted proteins, or internal structure that can act as a stop signal [31–34]. More broadly, mechanical resistance near the engaged SP terminal can be overcome by repetitive force application, however, unfolding rates are strongly dependent on the type of secondary structure present and the strength and extent of its connectivity with the SP core (such as van der Waals vs. hydrogen bonding, buried vs. solvent-exposed structure). A specific factor affecting the mechanical resistance is the anisotropy of the protein structure, which inherently gives rise to distinct responses when the direction of force application is varied. This aspect is particularly highlighted in proteins that include *β*-sheet structure given the asymmetric requirements involved in shearing and unzipping mechanisms. Shearing is achieved through nearly simultaneous removal of multiple inter-strand hydrogen bonds, therefore large mechanical forces are required, whereas unzipping involves sequential removal of these bonds with significantly smaller forces being required. Single-molecule force spectroscopy investigations of proteins containing *β*-sheet structure, such as the green fluorescent protein, ubiquitin, calmodulin or src SH3, reveal dramatically different unfolding resistance when forces are applied along the direction of N and C termini compared with directions that involve internal sites of the polypeptide chain [35–39].

In this paper, we focus on the action of two classes of AAA+ machines that process substrates at opposite ends of the length scale. ClpY performs unfolding and translocation of globular SPs with sizes smaller than or of the order of the machine, whereas the microtubule-severing enzyme spastin is responsible for the severing of MT assemblies, the longest and stiffest filaments in the cell, that dwarf the machine. Remarkably, such machines have the ability to remodel substrates of a range of sizes, such as the versatile ClpB/Hsp104 disaggregases that can disassemble large amorphous or fibrillar aggregates, but can be repurposed to perform degradation of globular proteins [40]. How do these machines adjust their action to this wide range of length scales while their available mechanical energy is limited by the capacity to hydrolyze ATP? It is likely that they accomplish spectacular unfolding of diverse substrates by taking advantage of the mechanical anisotropy in proteins. For example, degradation of tandem titin I27 domains mediated by the ClpX ATPase reveals branched pathways corresponding to the release of partially degraded fragments [41]. As indicated by simulations studies, the incomplete degradation process can be rationalized through lower barriers to unfolding single domain substrates, which may be oriented at the Clp ATPase surface to identify weak mechanical directions, and higher barriers to unfolding of multi-domain substrates due to the hindrance to rotational diffusion presented by the extra load [41]. Thus, it is reasonable to hypothesize that the relative orientation of the machine and the SP controls the unfolding process. In accord with the extreme length scales of SPs considered, we probe this hypothesis by performing computer simulations of variants of DHFR with distinct N-C terminals that dynamically reorient on the surface of the ClpY ATPase and of the microtubule-severing machines diffusing on the MT lattice. An important issue for understanding the action of AAA+ machines is the minimal assembly required to support the unfoldase function. In the Clp ATPase family, this is recognized to be the hexameric ring formed by the core AAA domains alone, therefore our simulations comprise the truncated ClpYΔI variant that lacks the auxiliary I domains. Such clarity is absent in MT severing, as several experimental studies [25,27,28] strongly suggest that additional domains are needed for severing. To account for the two possible scenarios, we adopt a twofold approach: in one set of simulations we consider solely the AAA ring (comprising NBD and HBD domains of each protomer), and in a second set the complete spastin machine (MIT, linker, and AAA region). The scope of this paper is to probe the SP unfolding action of these machines, therefore, in the spastin case, we focus exclusively on the proposed “unfoldase” severing mechanism, according to which the motors pull away subunits from a MT filament by using the mechanical work of sequential ATP hydrolysis seen in the majority of proteins from the AAA+ family [29,42].

Our results indicate that the critical breaking force for the removal of a MT fragment is at its minimum when the full spastin machine is present (the motor domain, the MIT domains, and the connecting linkers) and the interactions between the MIT domains and the surface of the MT lattice are in the interval 1.0 to 2.5 kcal/mol. By contrast, the use of only the motor domains or of the full machine with fixed MIT domains, results in increases in the breaking force by 100% or, respectively, 35% compared to the optimal scenario. Moreover, we found that the lowest value of the breaking force in our current simulations is comparable to the force yielded by our previous studies of the role of pulling in bending and breaking MT filaments where the filament could orient freely in space [43], which indicates that the full spastin machine can take advantage of the direction of least resistance for the disassembly of a MT lattice. We also find that unfolding of globular SPs mediated by the Clp ATPase is strongly modulated by the local DHFR interface initially engaged by the machine and the orientation of the SP at the lumen of the Clp pore. In both N- and C-terminal pulling of the wild-type DHFR, strong mechanical resistance of the *β*-sheet yields branched pathways that correspond to two SP orientations. In one case, the Clp-mediated pulling is applied in the direction nearly parallel to the *β*-sheet registry and unfolding is effected through a “shearing” mechanism that corresponds to a high-energy barrier. In the second case, pulling is applied nearly perpendicular to the *β*-sheet registry allowing unfolding to proceed through an “unzipping” mechanism with an associated lower-energy barrier. By contrast, unfolding and translocation of circular permutant (CP) variants of DHFR do not involve large barriers upon initial SP engagement, but mechanical resistance associated with internal structure can hinder these processes and result in long dwell times.

## 2. Materials and Methods

### 2.1. Homology model for the spastin machine

In our simulations we used two models for a spastin machine: (i) only the AAA motor domain in each protomer, and (ii) the full spastin machine, consisting of the motor region, the microtubule-interacting and trafficking (MIT) domains, and the flexible linkers that connect the motor domain with the MIT domain in each protomer. The main function of the MIT domain is to interact with the MT surface and thus to facilitate the placement of the AAA+ motor assembly on the MT lattice [44]. For (ii) we built a spastin hexamer model for which we attached the MIT domains to the N terminal end of the NBD domains of each protomer through flexible linkers. The protein sequence for the MIT domain and linkers was submitted to the HHPRED web server [45] to identify spastin’s MIT domain using a multi-template approach. Then using Modeller (version 9.23) [46], we built a homology model of the whole spastin machine based on the 6P07 (spastin + E15 peptide) and the 3EAB (spastin MIT domain) Protein Data Bank (PDB) structures, along with linkers (~60 amino acids). The obtained spastin hexamer complex model was then converted to a coarse-grained (CG) model by extracting the C*α* atoms from the complex. Finally, we mounted the CG model of the complex on a MT lattice with 8 dimers per protofilament (MT8) using Pymol [47].

### 2.2. Coarse-grained Model for the Unfoldase Action of Spastin on Microtubules

For the action of spastin on MTs, all simulations were performed using the self-organized polymer (SOP) model accelerated on GPUs (gSOP version 2.0) [48,49]. The model uses the equation shown below to determine the different interactions within the protein, described by the total potential (*V_T_*), that will dictate the dynamic behavior of the structure in time. The finite extensible nonlinear elastic (*V_FENE_*) potential represents the backbone of the structure, the full Lennard-Jones potential 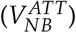 represents the native non-bonded interactions in the structure, while the repulsive Lennard-Jones potential 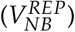 represents the non-native non-bonded interactions in the structure:

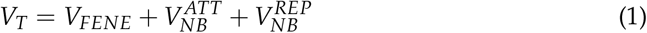

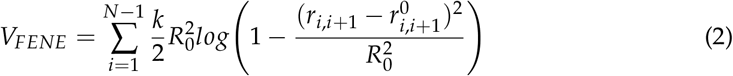

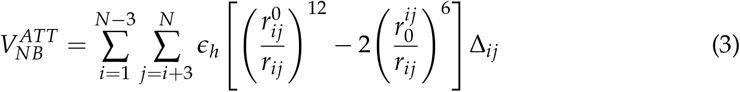

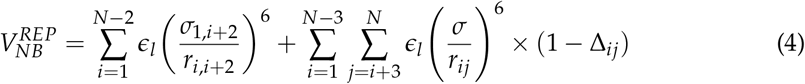

The strength of contacts for the non-bonded native interactions is given by the parameter *ϵ_h_* in Eq.3, which changes based on the lattice model used. The values for the contact strength (*ϵ_h_*) for each MT model used in our simulations are listed in Table. 1. The remaining parameters are: the frictional coefficient (*ζ*) set to 50, the spring constant (k) for covalent interactions set to 20.0 kcal/(mol ·Å^2^), *R*_0_ = 2.0 Å, and *r_i_* = 3.8 Å, for 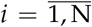, where N is the total number of residues. *r_i,j_* represents the distance between two residues, *i* and *j*, while 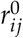 is its value in the native structure. The other parameters specifically used in pulling simulations are the cantilever spring constant, *k_trans_* = 0.025 kcal/(mol ·Å^2^), and the displacement of the cantilever during a simulation, Δ*x* = 0.0008 Å. The pulling speed (*v_f_*) is calculated using 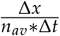, where *n_av_* is the number of steps and Δ*t* is the integration time step of 40 ps. In the cryo-EM hexameric structure of spastin in spiral conformation (PDB ID: 6P07), the E15 peptide chain is bound to the central pore loops of the spastin hexamer. We used the structure of the spastin hexamer, which we mounted on MT lattices, corresponding to a GDP configuration [50], of different sizes by attaching the N-terminal end of the E15 peptide to the C-terminal end of a *β*-tubulin monomer located centrally in the MT lattice. Then we performed a constant vector pulling along the direction of the E15 peptide oriented from the face A (facing the MT) to the face B (facing away from the MT) of the spastin hexamer.

**Table 1.** *ϵ_h_* (kcal/mol) values for intra-dimer, longitudinal, lateral and at the seam of the MT lattice, for spastin, and for interactions between spastin and the MT lattice.

|  |  |  |  |
| --- | --- | --- | --- |
| MT Intra-dimer | 1.9 | Intra-protomer | 1.3 |
| MT Longitudinal | 1.0 | Inter- protomer | 1.3 |
| MT Lateral | 0.9 | MT-Protomer | 1.0 |
| MT Seam | 0.9 | Protomer-E15 | 1.0 |

### 2.3. Simulation setups for the Unfoldase Action of Spastin on Microtubules

We performed simulations using a 3×3 dimers long MT lattice fragment (MT3 × 3) to explore the breaking of *αβ*-tubulin hetero-dimers (PDB ID: 1JFF) from the lattice fragment under the proposed unfoldase action of the spastin ATPase hexamer. At least two residues on the lumen side of each end monomer of the MT fragment were fixed to hold it in place, as shown in Fig 1A and 1B. We mounted the spastin AAA+ motor (Motor) in the spiral conformation on this MT fragment with the E15 peptide, covalently linked to the CTT from a central *β*-tubulin monomer, bound inside the central pore of the spastin hexamer (Fig. 1C). We fixed selected N-terminal residues from protomers in the spastin motor to hold it on the surface of the MT fragment, as otherwise the motor would move away from the MT along with the unfolded CTT, as soon as force is applied to the CTT. We performed pulling simulations using this system to identify the orientation of the motor on the lattice that leads to the most efficient breaking of fragments from the MT. Based on the results from this study, next we created a complex system consisting of a spastin’s AAA+ hexameric motor connected to the 6 MIT domains through the flexible linkers (HEX-MIT), as described above. We mounted the HEX-MIT on a 8 dimers long and 13 protofilament MT lattice (MT8) with the E15 peptide, covalently linked to the CTT from a specific *β*-tubulin monomer, bound inside the central pore of the motor in the orientation from the cryo-EM structure, as shown in Fig 1D.

**Figure 1.**
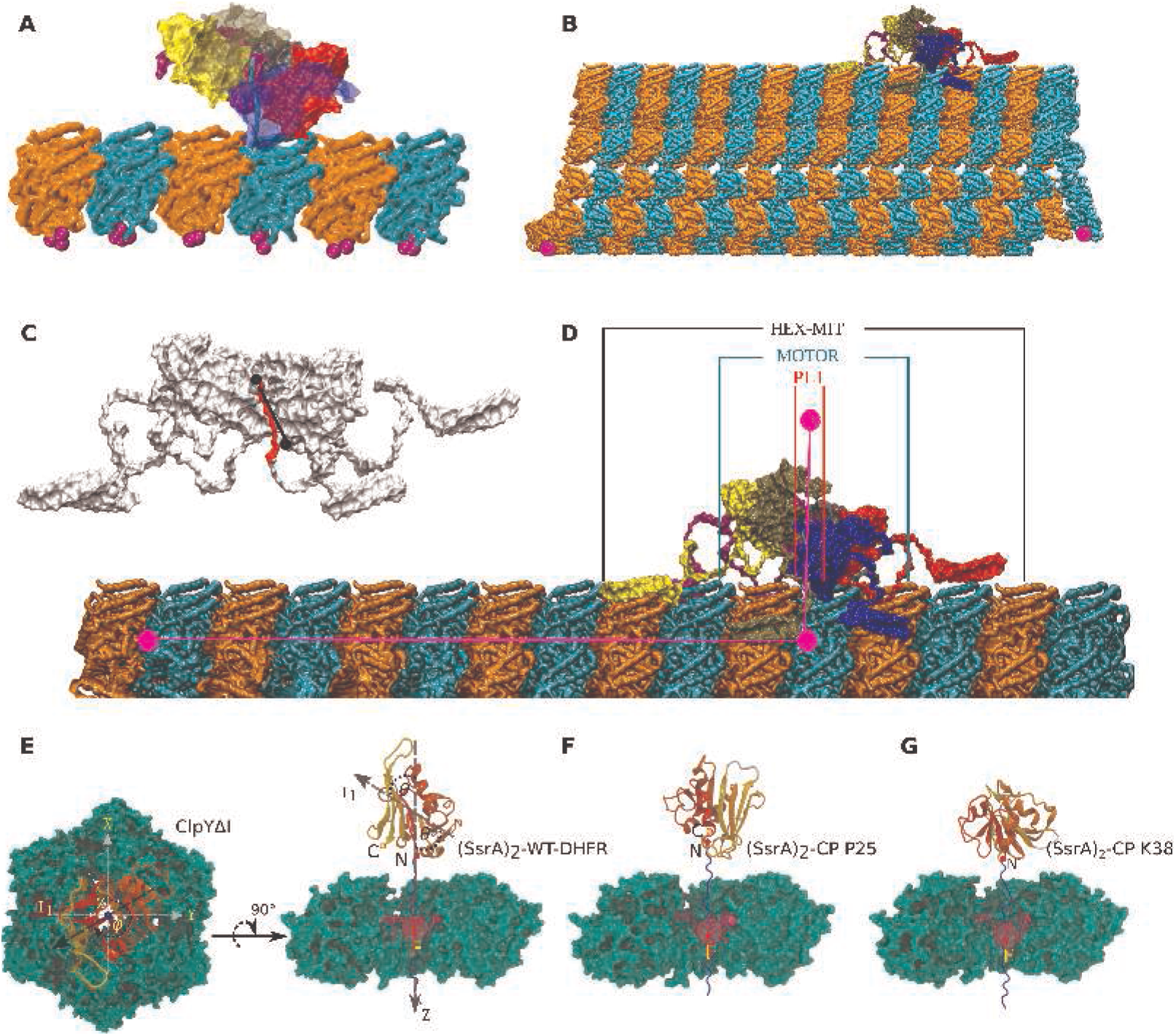
Configurations for the spastin and ClpY machines and their substrate proteins (SP) probed in our simulations. (A) side view of the spastin motor mounted on a MT3 × 3 fragment (*α* tubulin: orange, *β* tubulin: cyan, fixed residues: magenta); (B) side view of the entire spastin machine mounted on a 8 dimers long, 13 PF MT lattice; (C) cut-out side-view of the CTT (red) bound in the central pore of the spastin hexamer; (D) side view of the spastin machine mounted on a PF showing the various angles calculated in our pulling simulations: between the main axis of the PF (pink) and the principal axis of symmetry of the spastin machine (HEX-MIT), of the spastin motor only (Motor), or of the pore loops 1 (PL1) only; (E) Top and side view of the ClpYΔI (green)-SP system with polar angle *θ* and azimuthal angle *ϕ* indicated. SP is a fusion protein comprising the unfolded (SsrA)2 peptide (blue) and the DHFR domain (color-coded according to secondary structure). ClpY pore loops (red) are also indicated. (F)-(G) Circular permutant variants of DHFR with engineered N- and C-terminals at the (F) P25 and (G) K38 positions are also considered.

For the MT3 × 3 complex we used different set-ups characterized by fixing the N-terminal residues of various spastin protomers, which are in contact with the MT lattice: (i) fixing the N-term position for all the 6 protomers (chains A to F), and (ii) fixing the N-term residue for only two protomers (chains A and E). For the MT8 lattice, to hold it in place, we fixed one residue each on the first and last dimer thus fixing positions on both the minus and the plus end of the lattice, respectively. Then we probed different set-ups corresponding to either fixing residues from the MIT domains or altering the interaction strength (*ϵ_h_*) between the spastin motor, the MIT domains, the CTT and the MT lattice: (i) fixed all MIT domains on the MT lattice, with the contacts (*ϵ_h_* = 1.0 kcal/mol) defined between MT-MIT and Motor-*β*-CTT listed in Table 1 [Protomer-E15]; (ii) fixed MIT domains on the MT lattice, with the contacts (*ϵ_h_* = 1.0 *kcal*/*mol*) defined between the E15 peptide of the MT and the PL of the severing enzyme; (iii) fixed MIT domains of consecutive chains (A and B) and opposite chains ((B and E) with *ϵ_h_* = 1.0 kcal/mol between MT and MIT domains; and (iv) free MIT domains with the *ϵ_h_* values ranging from 1.0 to 4.0 kcal/mol between MT and MIT domains. To mimic the proposed unfoldase action of spastin, we performed our simulations by pulling on the C-terminal end of the E15 peptide, as described in the text, at a speed of 2 *μm/s*, using a regular lattice model for the MT filament, which represents the GDP type lattice, as discussed in our previous work [50]. We performed 3 independent trajectories for each set-up.

### 2.4. Data analysis for the Unfoldase Action of Spastin on Microtubules

For each type of simulation, we monitored the breaking patterns of the MT lattice, the breaking of spastin hexamer into lower order oligomers, and the loss of contacts between the spastin machine and the MT lattice. We identified the orientation of spastin with respect to the MT lattice during the severing process, by calculating three angles using Visual Molecular Dynamics [51]. The angle *θ* is the angle between the long axis of the pulled protofilament (PF), obtained between the center of mass of the dimer on the plus end and the center of mass (COM) of the dimer on which the pulled CTT is located, and the main principal axis of the full spastin hexamer, including the linker and the MIT domain (Hex-MIT), as shown in Fig 1D. This angle provides information about how the severing enzyme reorients/behaves upon fixing the MIT domain(s) or varying the interaction strength of the MIT domain with the MT lattice. The angle *ϕ* is the angle between the PF where the pulled CTT is located and the main principal axis of the spastin motor only (motor). This angle characterizes the orientation of the motor on the lattice as a result of the relative motions of the NBD and HBD domains, and/of the protomers. Finally, the angle *ψ* is the angle between the long axis of the pulled PF and the axis of the pore loop 1 (PL1), defined as the unit vector connecting the C*α* atoms of the highly conserved pore loop residues K555 from chains A and F of the spastin hexamer, which interact directly with the CTT, as shown in the Figure 1C and 1D. We also calculated the fractional loss of native contacts (*Q_N_*) for the pulled protofilament (PF6) of the MT lattice and categorized them into longitudinal (along the PF) and lateral (adjacent to the PF: East and West interface) contacts, following our earlier work [43,52], as detailed below. Based on the above angles and the *Q_N_* values, we obtained the free energy landscapes corresponding to the orientation of the severing enzyme on the lattice.

### 2.5. Implicit Solvent Model of ClpYΔI with DHFR or its Circular Permutation (CP)

Simulations of DHFR remodeling assisted by ClpY are performed using the EEF1 (Effective Energy Function) implicit solvent model [53,54], which ensures computational efficiency in probing interactions at atomistic resolution.

Wild-type and CP CHFR variants share the same three–dimensional structure, which is modeled using the crystal structure of the *Escherichia coli* protein with PDB ID 5W3Q, but have different polypeptide terminals. In CP variants, new terminals of the polypeptide chain are engineered through cleavage of the C–N peptide bond between sequence positions 24-25 and 37-38, respectively, with new N terminals being located at the P25 and K38 sites. To ensure chain connectivity, CP variants include a (Gly)_5_ linker that connects the wild-type N and C terminals. We obtained low-energy configurations of the initially extended structure of the (Gly)_5_ linker by performing energy minimization of the DHFR domains, which comprised 1000 steps using the steepest descent (SD) algorithm and 1000 steps using the adopted-basis Newton-Raphson (ABNR) method. During the energy minimization steps, constraints were applied on DHFR atoms, except for those of the linker residues, to maintain them at fixed positions corresponding to those in the crystal structure. An (SsrA)_2_ degradation tag (SsrA sequence AANDENYALAA) was covalently attached at the N- or C-terminal of each folded domain to initiate translocation in the N-C and C-N direction, respectively, of the resulting fusion protein through the ClpYΔI nanomachine.

The center of mass of the ClpY ATPase is maintained near the origin of a Cartesian reference system and the ClpY pore axis is aligned with the *z*–axis, which is oriented such that the *cis* (proximal) side corresponds to *z* < 0 and the *trans* (distal) side to *z* > 0 (Figure 1). The SP is initially oriented such that its principal axis of inertia is aligned with the *z*–axis and its center of mass is located at *z* ≃ −50 Å on the *cis* side of ClpYΔI ATPase. The (SsrA)_2_ peptide, which has an extended conformation, is partially inserted into the ClpY pore so that the SP can be firmly engaged by the nanomachine. Distinct initial configurations are generated for each simulation trajectory by rotating the SP through an arbitrary azimuthal angle about the *z*–axis. We use the CHARMM molecular modeling package [55] to perform Langevin dynamics simulations at T = 300 K, with a friction coefficient of 5 ps^-1^ and a time step of 2 fs. Simulations were performed on Extreme Science and Engineering Discovery Environment (XSEDE) supercomputer resources [56].

### 2.6. ClpYΔI Allosteric Motions

Our computational model describes the interaction between ClpY and SP through stochastic binding and release events. To this end, the allosteric cycle of the Clp ATPase is modeled using sequential single-protomer conformational transitions between states of high and low SP binding–affinity that correspond to “open” and “closed” pore configurations. Crystallographic structures of these two configurations are obtained from *Escherichia coli* ClpY, namely PDB IDs 1DO2 (open pore) and 1DO0 (closed pore) [6]. We consider the truncated ClpYΔ*I* variant that excludes specific auxiliary I–domains (residues 111–242) in order to facilitate the understanding of the common mechanisms of Clp ATPases. To account for the absence of the I domain, each ClpY protomer is modeled using two polypeptide chains.

Conformational transitions of each protomer are described using the targeted molecular dynamics (TMD) approach [57], which probes conformational transitions between two configurations of the molecule by minimizing the root mean square deviation (RMSD) between them. Each hemicycle (open → close or close → open) of hexameric ClpY incorporates six sequential transitions of individual protomers. The subunit undergoing the first transition in the hemicycle is randomly selected and subsequent transitions follow the clockwise ring order as viewed from the *cis* side of ClpY. In each step, the centers of mass of all protomers except for the active protomer are constrained to their current position. Each cycle has a total duration *τ* = 120 ps and the effective pulling speed is 1 Å/ps given the 10 Å excursion of pore loops. Our simulations of ClpY–mediated unfolding and translocation of I27 SPs indicate that results obtained using this effective speed are in agreement with those obtained at lower speeds (0.2 Å/ps and 0.02 Å/ps) [41] and with atomistic simulations of bulk mechanical unfolding [58]. It is also important to emphasize that the time scales probed in implicit solvent simulations correspond to longer effective biological times [59–61].

### 2.7. External Repetitive Force Coupled with Allosteric Motions

An external repetitive force is applied along the pore axis, during the open →close hemicycle, on the backbone heavy atoms of the SP that are transiently located within the central pore of the ClpY ATPase in order to accelerate the SP unfolding and translocation process. The relevant heavy atoms of the SP are identified according to |*z* − 〈*z*_loop_〉| < 5 Å, where *z* represents the axial coordinate of the atom and 〈*z*_loop_〉 represents the average *z* coordinate of Tyr91 amino acids of central channel loops of ClpY at the beginning of each cycle. The magnitude of the force is obtained from a Gaussian distribution with a mean value that reflects the mechanical resistance of the SP and the force is uniformly distributed onto the backbone heavy atoms of the SP. For the wild-type DHFR, the average external force is 700 pN for N-C translocation and 400 pN for C-N translocation, with standard deviations of 50 and 20 pN, respectivel; whereas, for the CP variants, the average external force is 400 pN for P25 and 600 pN for K38, with standard deviation of 20 pN (3). The number of backbone atoms that are instantaneously located in the ClpY pore region is about 20 atoms, therefore the applied force per atom has an average of ≃ 20 pN for WT-DHFR in the C-N direction, ≃ 35 pN for WT-DHFR in the N-C direction, ≃ 20 pN for CP P25 in the N-C direction and ≃ 30 pN for CP K38 in the N-C direction.

### 2.8. Fraction of Native and Non–Native Contacts

The fraction of native contacts (Q_N_) is computed as 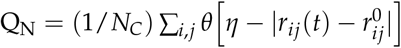 where N_C_ is the number of native contacts, *r_ij_*(*t*) is the distance, at time *t*, between residues *i* and *j* and 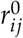 is the corresponding native distance. *θ*(*x*) is the Heaviside step function for which *θ*(*x*) = 1 if *x* ⩾ 0 and *θ*(*x*) = 0 if x < 0, and the tolerance is *η* = 2 Å. For MTs, N_C_ represents the number of native inter-subunit contacts formed between the pulled tubulin subunit and its MT lattice neighbors within the PF. The cutoff distance for residue-residue interactions in the native configuration is set to 13 Å. For the DHFR domains, N_C_ represents the number of native intra-domain contacts, with *r_ij_*(*t*) identified as the minimum distance between any two heavy atoms of residues *i* and *j*. Here, the sequence separation between residues must be larger than 2, i.e. |*i* − *j*| > 2, and the cutoff distance is set to 6 Å. The fraction of non–native contacts (f_NN_) is defined as f_NN_ = N_NC_/N_C_, where N_NC_ is the number of non–native residue pairs identified using the 6 Å cutoff.

### 2.9. Translocated Fraction and Waiting time of the polypeptide chain

Th translocated fraction, x(t), is defined as the instantaneous fraction of amino acids that have progressed to the *trans* side of the ClpY pore, i.e. the axial location of the *C_α_* atom lies beyond the maximum excursion of the ClpY pore loops, *z_i_*(*t*) > *z_trans_* = 12 Å. The translocation line, *I*(*t*), is determined by using the sequence position of the most recently translocated amino acid, with *Z_I_*(*t*) = min{*z_i,trans_*(*t*)}.

To describe the translocation hindrance per residue at the ClpY pore lumen we determine the so-called waiting time [62], *w*(*I*), which is estimated based on the residence time of residue I in the vicinity of the ClpY pore entrance such that *Z_I_*(*t*) ≳ *Z_lumen_* = −8 Å.

### 2.10. SP orientation near the ClpYΔI Pore Surface

To capture the orientation of the folded fragment of the SP with respect to the ClpYΔI pore axis, we use the angular degrees of freedom in the spherical coordinate system. The polar angle *θ* represents the angle between the first principal axis of the SP and the z-axis. The azimuthal angle *ϕ* represents the angle between the projections of the principal axis of SP and of the position vector of the center-of-mass of one subunit of ClpYΔI onto the plane perpendicular to the z-axis.

## 3. Results and Discussion

### 3.0.1. Spastin machine without MIT domains mounted on a MT3 × 3 lattice

In order to probe spastin’s orientation on a MT lattice during the severing process and the implications for the action of the machine, we started by investigating the behavior of the spastin motor alone. As described in the Methods, we mounted the spastin motor’s face A onto a 3 × 3 dimers-long (MT3 × 3) lattice with the CTT of the central *β*-tubulin monomer bound inside the pore of the motor, as found in the recently solved cryoEM structures of spastin [25]. For these simulations, we used two set-ups corresponding to fixing the N-terminal residues (1) of all the motor protomers, and (2) of only two protomers opposite to each other (from chains A and E). To mimic the mechanical action associated with the proposed unfoldase model of severing [30], we applied a constant loading force, at a pulling speed of 2 *μm*/*s*, to the C-terminal residue of the central *β*-tubulin monomer. This approach mimics the set-up from LOT experiments and simulation studies for other AAA+ machines. We note that more detailed models, such as the approach based on targeted transitions between spastin motor’s configurations [63] while acting on its substrate, which we employed for the ClpY simulations, are prohibitive due to the large size of the substrate (the MT lattice). The first event observed during these simulations in both set-ups was the unfolding of the C-terminal region of the pulled *β*-tubulin corresponding to the unraveling of its H11, H11’, and H12 helices and the E10-strand. Fixing the ends of all the protomers on the MT, i.e., using set-up (1), results in the unfolding of these regions at the first (FBF), which is also the critical (CBF), breaking force of 750 pN and leads to the loss of contacts between the pore-bound CTT and the PLs of the spastin motor, ending with the exit of the CTT through face B of the motor (Fig. S1A). The unfolding of the pulled *β*-monomer continued up to its E5-strand [64]. In contrast, when only the N-terminal ends of chains A and E in the motor were fixed, the FBF was lower (670 pN) than in set-up (1) and corresponded to the unfolding of the pulled *β*-monomer with the CTT remaining bound inside the pore of the motor (Fig S1B). While in set-up (2) there was no additional unraveling of the pulled *β*-monomer, we found that continuous pulling on its CTT results in the unfolding of the N-terminal domains of the two spastin motor protomers with ends fixed, along with the stretching of the unfolded C-terminal region of the pulled *β*-tubulin. These events led to the lifting of the motor upwards from the MT surface, while remaining oriented parallel to the MT surface, followed by the CTT losing contacts with the PL residues and exiting the pore of the spastin motor. Next, both set-ups led to the same event: the loss of lateral and longitudinal contacts between the pulled *β*-tubulin monomer and the rest of the MT fragment. This resulted in the *β*-monomer in the first set-up, and the pulled dimer in the second set-up moving closer to the gate between the end protomers of the spastin motor. Finally, we saw the loss of the respective monomer or dimer tubulin from the lattice. Thus the net result of these types of simulations is the extraction, from the MT lattice, of one substantially unfolded tubulin monomer in the first and, respectively, a dimer in the second set-up.

In summary, these simulations, involving only the spastin motor and the MT3 × 3 fragment, led to either the unfolding of most of the pulled monomer, when all spastin protomers were fixed, or to the unfolding of the fixed protomers’ N-terminal ends (residues range 455-512: 57 residues long), along with the C-terminal of the pulled monomer, when fixing the N-terminal of only protomers A and E. Importantly, in both set-ups the spastin motor maintained its original (parallel) orientation with respect to the surface of the MT fragment throughout the simulations. Based on experimental data, [25,65] none of these events are plausible results of the severing action, as they involve extensive unfolding of a tubulin subunit and of the spastin motor. Nevertheless, we note that the unfolding of the N-terminal domains of the fixed protomers had the beneficial effect of allowing the motor enough freedom to adjust its position with respect to the surface of the MT3 × 3 lattice and to the direction of the CTT peptide. This in turn enabled the PL’s continuous grip on the tubulin peptide fragment found inside the spastin motor for the duration of the trajectory. Our observations show that, to function properly, spastin needs to have the ability to change orientation such that the motor can closely track the orientation of the tubulin chain to be pulled. This in turn would allow the PLs from its central pore to exert a constant grip on the substrate. Because in our motor-only simulations this tracking ability came at the expense of the unfolding of regions from the protomers in the motor, we concluded that the spastin motor cannot sever MTs by itself. This finding gives molecular support for similar proposals from the literature [26,66] and provides insight into the rationale for the use of more than just the motor region to induce MT severing.

### 3.0.2. Spastin machine with fixed MIT domains mounted on a MT8 lattice

Our results from above show that, for optimal processing of the substrate, the severing protein needs to have enough free space to adjust its orientation with respect to the MT filament, while still being located close enough to the MT to induce its severing. The gain of free space during severing, which we saw above as the result of the unfolding of parts from the motor domain, can alternatively be envisioned as the stretching/unfolding of a long flexible chain, with no defined tertiary structure. This suggests a plausible functional role for the long flexible linkers that connect the motor domain of spastin to its MIT domains. Thus, next we carried out simulations where we modeled the spastin machine consisting of the motor, the MIT domains, and the long (~ 60 residues) flexible linkers (HEX-MIT) that connect them. We note that the length of the linker is similar to the length of the N-terminal portion of the protomers that unfold in our MT3 × 3 simulations with fixed A and E domains. For this set of simulations, we modeled the spastin machine mounted on an 8 dimers long, 13PF MT lattice (MT8), as shown in Fig. 1B. In addition, we fixed the N-terminal residue of each protomer’s MIT domain (fixed MIT). We carried out two simulation set-ups: for the first set-up we used an interaction strength (*ϵ_h_*) of 1 kcal/mol between the MT and the MIT domains, and between the MT (including the CTT) and the entire spastin motor. For the second set-up, we kept the interaction strength between MT and MIT the same as in set-up one, while reducing the interaction between the spastin motor and MT to only the interactions between the CTT and the PLs (*ϵ_h_* = 1.0 kcal/mol). In the starting configuration, the principal axes of the HEX-MIT, the motor, and the central pore are all perpendicular to the long axis of the pulled PF. Similar to the above simulations, we applied a constant loading rate pulling force at the C-terminal end of the CTT, as detailed in the Methods section. The first event in both set-ups was the unfolding of the C-terminal domain (helices H12 and H11) of the pulled *β*-tubulin monomer at a 100-200 pN force, which coincides with the first event in the MT3 × 3 simulations. In these new set-ups the motor moves up away from the MT lattice along with the pulled CTT, while the fixed MIT domains hold the machine on the lattice. This was similar to what we observed in the fixed N-ter of chains A and E on MT3 × 3 set-up discussed above. The unfolding of the *β* monomer continued as the CTT got threaded through the central pore, lost contacts with the PLs, and exited the motor under the action of a ~ 230 pN FBF, as shown in the Fig S6A and S6B. From this point on, the motor remained parallel to the longitudinal axis of the MT lattice, allowing for further unraveling of the pulled *β*-monomer (up to residue 372), and the loss of contacts between the pulled PF and its lateral neighboring PFs in the lattice. The CBF at ~ 430 pN corresponded to the loss of lateral contacts of the pulled PF, followed by the breaking of the northern longitudinal interface of the pulled dimer and the unzipping of the resulting PF fragment towards the minus end of the MT filament, finally resulting in the extraction of a 6 dimers fragment from the MT lattice.

During the simulations following the first set-up, the principal axis of the severing enzyme (*HEX_MIT_*) remains perpendicular to the long axis of the MT lattice for the duration of the run. In contrast, the motor moves up from the lattice and its principal axis switches from perpendicular to forming angles of ~ 10° - 40° with the long axis of the unzipped PF. The switch in the orientation of the motor (Fig. S2) was due to the relative motions and fluctuations in the protomers. Finally, the PL1s align along the direction of the pulled PF, as shown in Fig. S3. In the second set-up, the HEX-MIT makes angles of 50°-70° with the longitudinal axis of the pulled PF during the loss of the lateral interfaces and finally settles at lower angles after the hexamer dissociation. At the same time, the central pore remains perpendicular to the MT lattice for the duration of the simulations. The breaking of the hexamer into trimers can be observed in the PF vs PL plot from Fig.S3.

In summary, from this set of simulations we found that fixing all the MIT domains resulted in holding spastin on the lattice even after PF severing and motor dissociation had occurred, We therefore modified our simulation set-up by fixing only two MIT domains, either from consecutive protomers (chains A and B) or from opposite protomers (chains B and E). Keeping the interaction strength between MT and MIT domains set to 1.0 kcal/mol, we fixed the MIT domains of only the chains A and B and followed the pulling procedure from above. We found that all the free MIT domains, with the exception of the MIT domain in chain F, lost their contacts with the MT lattice at the start of a trajectory and fluctuated freely during the simulation. The MIT domain of chain F remained attached to the lattice until the force reached the FBF value of ~ 110 pN. This force was responsible for the unfolding of the pulled *β*-monomer from its C-terminal end up to the B10 strand (residues 374-429), resulting in the lifting of the motor above the lattice. During this first event, we observed a rapid switching in orientation of the principal axis of the full machine due to the paddling-like motion of the hexamer’s terminal protomers, as seen in the FEL and angles plot for PF vs HEX-MIT (Figs S4 and S5). We note that, in contrast, the motor and the PL1 loops maintain their original perpendicular orientation with respect to the lattice. The next event corresponded to the unfolding of the two fixed MIT domains along with the stretching of the C-terminal end f the pulled *β*-monomer until the CBF of ~ 450 pN is reached (Fig. S6C). Under the action of the CBF, the pulled *β*-monomer loses lateral contacts with monomers from the adjacent PFs and the plus end longitudinal interface of the pulled dimer breaks. Next, the CTT is released from the motor as a result of the loss of its contacts with the motor. This event is accompanied by the unzipping of the pulled PF towards the minus end of the MT lattice, which is in the opposite direction compared to the PF unzipping observed in the fixed MIT domains runs. The minus end oriented unzipping results in higher angles between the PF and the motor, with the PLs aligning along the unfolded tubulin. Finally, the motor rotates such that the unzipped PF moves away from the PLs, leading to the extraction of 6 tubulin dimers from the lattice.

Upon fixing the MIT domains of chains B and E (on diagonally opposite protomers), with the two MIT domains on either side of the fixed domains remaining free (chains C and D and A and F), the principal axis of the HEX-MIT oriented perpendicularly to the pulled PF and the motor moved up along the unfolded part of the pulled tubulin monomer during the initial C-terminal unfolding of the *β*-monomer. Further pulling at the C-terminal end resulted in the unfolding of the fixed MIT domains. At an average CBF of ~ 447 pN, the pulled dimer loses its lateral contacts, followed by the breaking of its longitudinal northern interface resulting in the formation of 6- and 2-dimers-long PF fragments (Fig. S6D). Next, the pulled CTT gets threaded through the PLs and exits the motor. The 6-dimers-long PF fragment unzipped towards the minus end of the lattice and eventually detached from the lattice. During the simulations, the principal axes (main axis of inertia) of the HEX-MIT, the motor, and the PL aligned at ±20°with respect to the longitudinal axis of the filament, indicating that the severing enzyme stays parallel to the MT lattice.

### 3.0.3. Spastin machine with free MIT domains mounted on a MT8 lattice

The above simulations showed that severing enzymes require the presence of both the MIT domains and the linkers to enable an optimal orientation of the motor in the proximity of the surface of the MT lattice during severing. Fixing the MIT domains onto the lattice, however, results in their unfolding along with the unfolding of the C-terminal end of the pulled *β*-tubulin, which is not supported by experimental findings. To address this issue, next we carried out simulations where we varied the interaction strength between the MIT domains and MT lattice, as opposed to fixing them on the lattice. The magnitude of the interaction strength between severing proteins and MT filaments is unknown due to difficulties in carrying out relevant experiments and due to the prohibitively large length and time scale of atomistic simulations that could yield such energy terms. A similar problem regarding the interactions between another MT associated protein, kinesin-1, and MTs was recently solved using a coarse-grained based method, which yielded calculated unbinding forces between kinesin motor domains and MTs, at increasing loading rates, for a range of interaction strengths. Next, the calculated forces were compared with experimental force values. Finally, the interaction strength of kinesin-1 binding to MTs at low and high affinity, depending on the nucleotide state of the kinesin motor, which led to the best agreement between the calculated and the experimental values, was selected [67]. Due to the lack of experimental unbinding force data for severing enzymes, we could not follow exactly the approach employed for the kinesin-MT interactions. In turn, we chose to vary the interaction strength (*ϵ_h_*) between the MIT domains and the MT and to compare the outcome of our loading rate based pulling simulations with experimental findings. Moreover, in these simulations we probed how changes in the strength of interactions affects the breaking pathways of the MT and the stability of the spastin oligomeric state. We explored interaction strengths in the range 1.00 - 4.00 kcal/mol, keeping the selected value consistent in all six MIT domains. We note that any values below 1.0 kcal/mol result in the rapid detachment of the spastin machine from the MT as soon as a pulling force is applied to the CTT. Similar to the above pulling simulations set-ups, we monitored the MT lattice breaking pattern, the orientation of the severing enzyme on the lattice, and the fraction of native contacts lost (*Q_N_*) over the course of the simulation (Figs. 2–3 and S7-S11). Details of the simulations are provided in Table 2.

**Figure 2.**
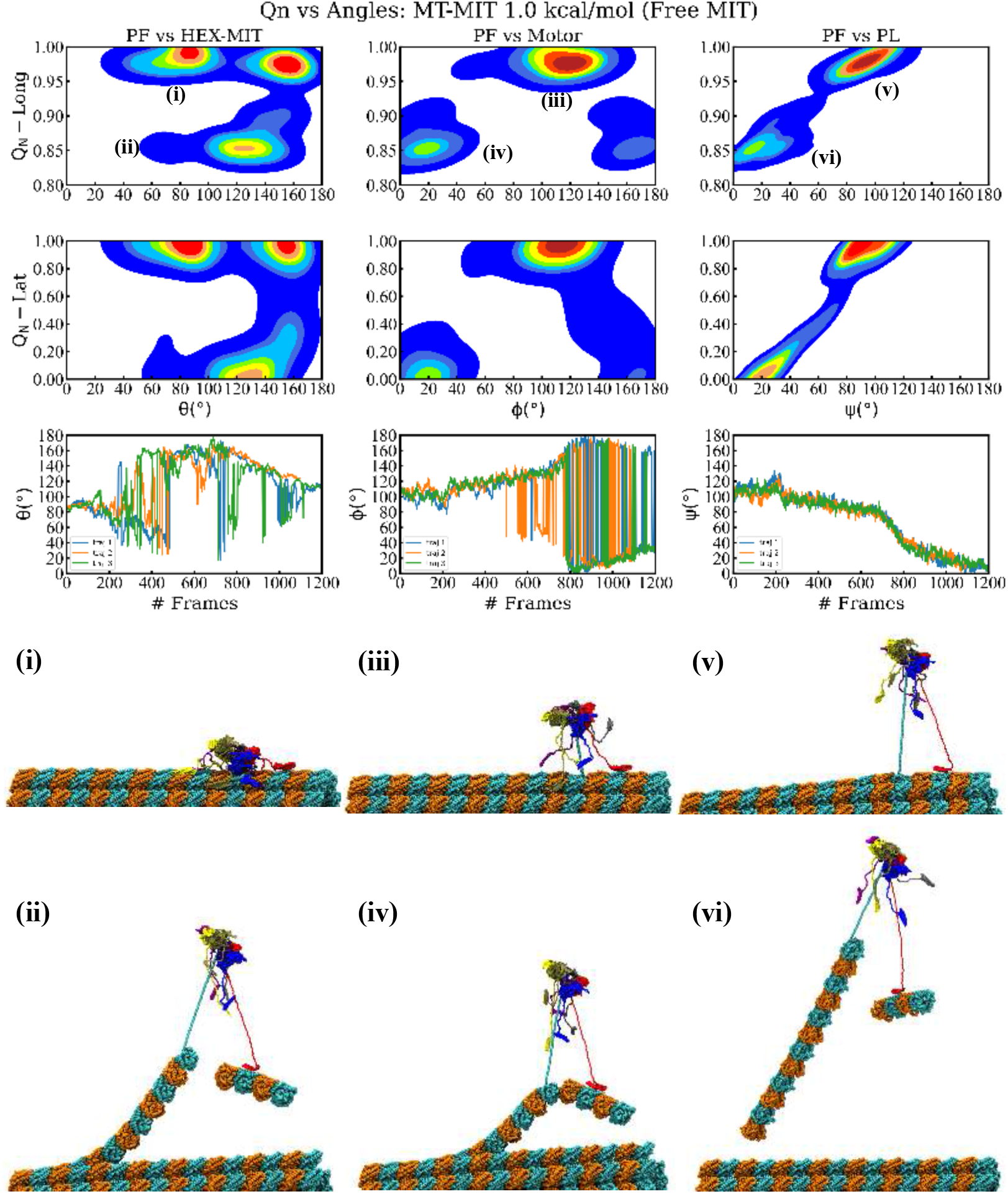
Results of the spastin machine acting on a MT filament for the interaction strength between its MIT domains and the MT lattice set to 1.0 kcal/mol. Top row plots show the free energy landscape in the plane of the fractional loss of longitudinal native contacts (*Q_N_*) of the pulled protofilament (PF6) and the angle made by the principal axis of the severing enzyme (HEX-MIT), of the motor, and of the central Pore loops (PL), respectively, versus the long axis of the pulled PF (PF). Middle row plots show the free energy landscape in the plane of the fractional loss of lateral native contacts of the pulled protofilament and the three angles from the top row panels. Lowest row plots depict the time evolution of the three angles from the upper plots versus the simulation frames. The representative structures corresponding to the labeled minima in the free energy plots are shown at the bottom.

**Figure 3.**
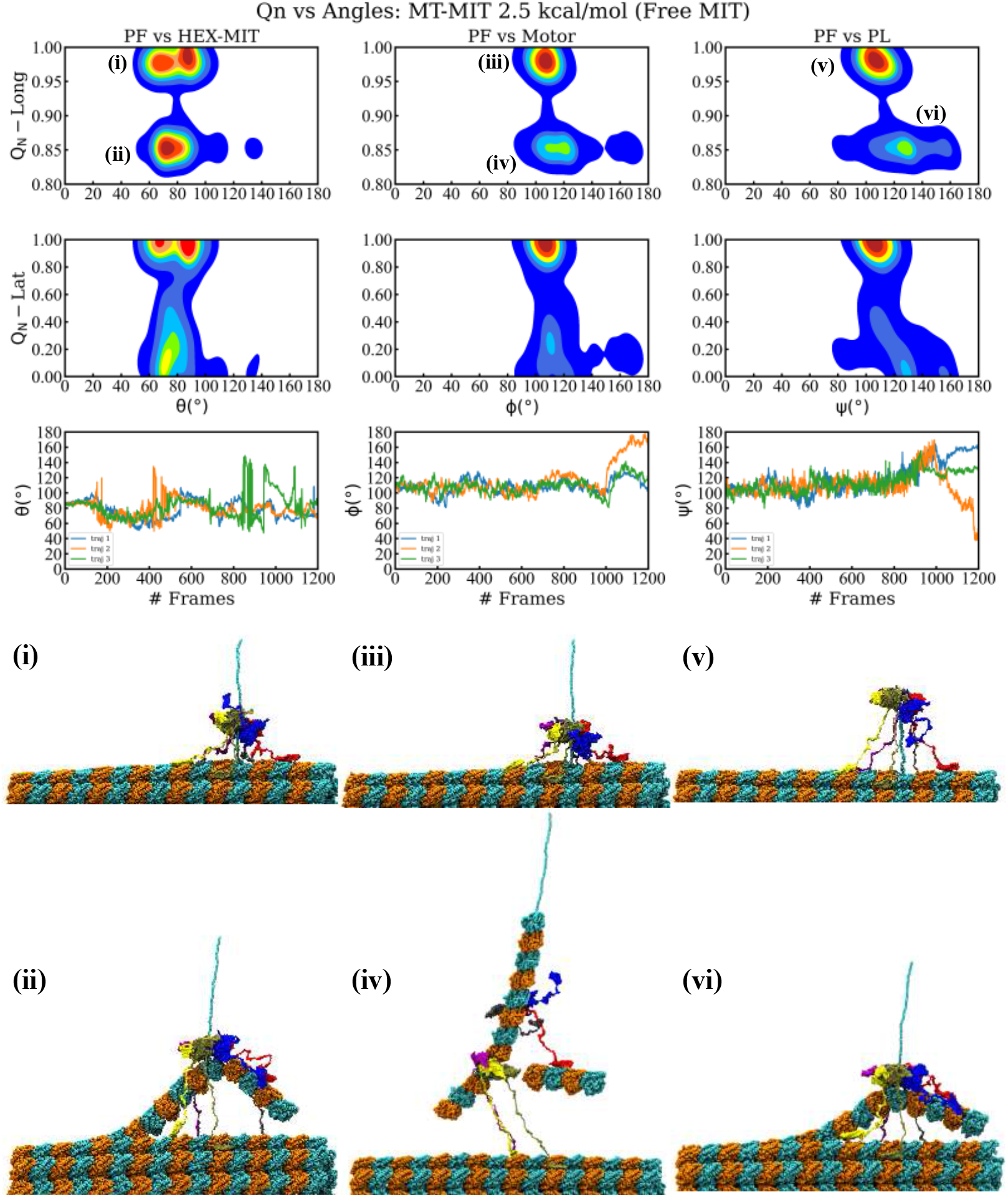
Results of the spastin machine acting on a MT filament for the interaction strength between its MIT domains and the MT lattice set to 2.5 kcal/mol. The plots follow the order detailed in Fig.2.

**Table 2.** Outcomes of simulations for the full spastin machine mounted on a MT lattice, with variable interaction strengths between the MIT domains in spastin and the MT surface.

| Set-up | $\epsilon_h$ (kcal/mol) | $\langle CBF \rangle$ (pN) | spastin oligomers | MT dimers lost (frequency) |
| --- | --- | --- | --- | --- |
| 1 | 1.00 | 380 | 0 | 6+2 (100%) |
| 1 | 1.00 | 380 | 0 | 6+2 (100%) |
| 1 | 1.00 | 380 | 0 | 6+2 (100%) |
| 2 | 1.50 | 345 | 0 | 6 (67%), 6+2 (33%) |
| 3 | 2.00 | 355 | 4+2 (67%), 3+3 (33%) | 6 (67%), 6+2 (33%) |
| 3 | 2.50 | 343 | 3+3, 0, 4+2 | 6+2 (67%), 6 (33%) |
| 4 | 3.00 | 373 | 3+3, 0, 4+2 | 6+2 (33%), 6 (67%) |
| 5 | 3.50 | 350 | 0 | 6 (100%) |
| 6 | 4.00 | 356 | 4+2, 0, 2+1+3 | 6+2 (33%), 6 (67%) |

We found that, as expected, a higher interaction strength leads to an increase in the number of MITs remaining in contact with the MT surface throughout the course of the simulation. For interaction strengths in the interval 1.00-1.50 kcal/mol only the MIT domain from chain B remains in contact with the MT, causing the severing motor’s principal axis to orient at ~ 120°, rather than being perpendicular, with respect to the long axis of the pulled PF due to the uneven anchoring of the spastin motor on the MT lattice (Figs. 2 and S7). Chain B remained attached to the pulled PF following the severing event, which resulted in the angle between the PF and HEX-MIT decreasing by ~ 40°after the loss of both longitudinal and lateral contacts. Importantly, at low interaction strengths, we found significant shifts in the orientation of the PL and of the motor in relation to the pulled PF due to the inherent mobility of the MIT domains, which allows the motor to follow the direction of the PF fragment that is eventually removed from the MT lattice. In these simulations, spastin detaches completely from the MT filament at the end of the severing event, as it remains anchored to the extracted PF fragment. Such trends diminished at higher interaction strengths. where more MIT domains maintain their contact with the MT surface for the duration of the run. Namely, at 2.00 kcal/mol (Fig. S8), three of the six MIT domains (from chains B, E, and F) remained in contact with the lattice, while at 2.50 kcal/mol (Fig. 3), a fourth MIT domain (from chain D) also adhered to the MT surface. The machine’s principal axis started perpendicular, but switched closer to parallel (angles ~ 20-30°) to the direction of the pulled PF at the end of the severing event, due to the more balanced anchoring provided by the fact that the MIT domains on opposite sides of the spastin motor preserved their contacts with the MT. In addition, this balanced anchoring led to only minimal adjustments in the orientation of the motor and of the central pore versus that of the pulled PF after the severing event, which can be seen in the FEL plots of the PL and motor. At the highest interaction strengths (3.00 and 4.00 kcal/mol), the MIT domains from all chains, except for chain A, remain bound to the lattice throughout the simulations. The severing machine began perpendicular to the lattice but had larger changes in orientation once the lateral and longitudinal PF contacts were lost due to the bound chains inducing the motor to lean towards one side, away from the freely fluctuating linker and the MIT domain of chain A (Figs. S10-S11). We again observed, similar to the simulations at intermediate values of the interaction strength, a lack of change in the orientation of the PL and the motor vs the direction of the pulled PF. Moreover, usually the MIT domains at higher interaction strengths keep the spastin machine in contact with the MT lattice at the end of the severing event, instead of the enzyme’s assumed behavior of dissociating from the lattice. Still, for the majority of the trajectories at 2.0 kcal/mol and at 3.0 kcal/mol, we found that the severing hexamer breaks into lower order oligomers (trimers or a tetramer and a dimer), which detach from the MT surface because they remain bound to the broken PF fragments. The modes of MT lattice breaking in our simulations are represented in the breaking pathways, which vary at all levels of interaction strength and are summarized in Fig. 4 and Table 2. All simulations began with the same initial unfolding of the pulled tubulin monomer’s C-terminal and led to the loss of lateral and longitudinal contacts in the pulled PF, as seen in the previous set-ups. The average CBF for the varying interaction strengths was 343-380 pN, which is the lowest value among all our different types of simulations. 57% of all trajectories resulted in the removal of a 6-dimer PF fragment, while the remaining 43% of runs resulted in a 6- and a 2-dimer fragment removed from the lattice. With the exception of 3.00 kcal/mol, all intervals of interaction strength showed a mixture between these two pathways. The most commonly followed pathway (38%) corresponds to the retention of the full spastin hexamer after severing a single 6-dimer PF fragment. Hexamer retention after losing the two PF fragments (19%) was the next likely pathway, followed by the pathway corresponding to the dissociation of the spastin hexamer into a tetramer and a dimer after cutting a single 6-dimers long PF fragment (14%). Tetramer/dimer (10%), two trimers (10%), and a trimer/ dimer/monomer (5%) formation were also identified pathways after severing the two PF fragments, and two trimers (5%) was the least found pathway for a single pulled PF fragment. The spastin motor was most likely to maintain its hexameric form at lower interaction strengths (1.00-1.50 kcal/mol), but we found it to be stable as a hexamer even at interactions strength as high as 4.00 kcal/mol. Starting at 2.00 kcal/mol, we found that the motor showed a tendency to dissociate into lower order oligomers. Interestingly, none of the trajectories using an interaction strength of 3.50 kcal/mol showed any dissociation of the spastin hexamer into lower order oligomers.

**Figure 4.**
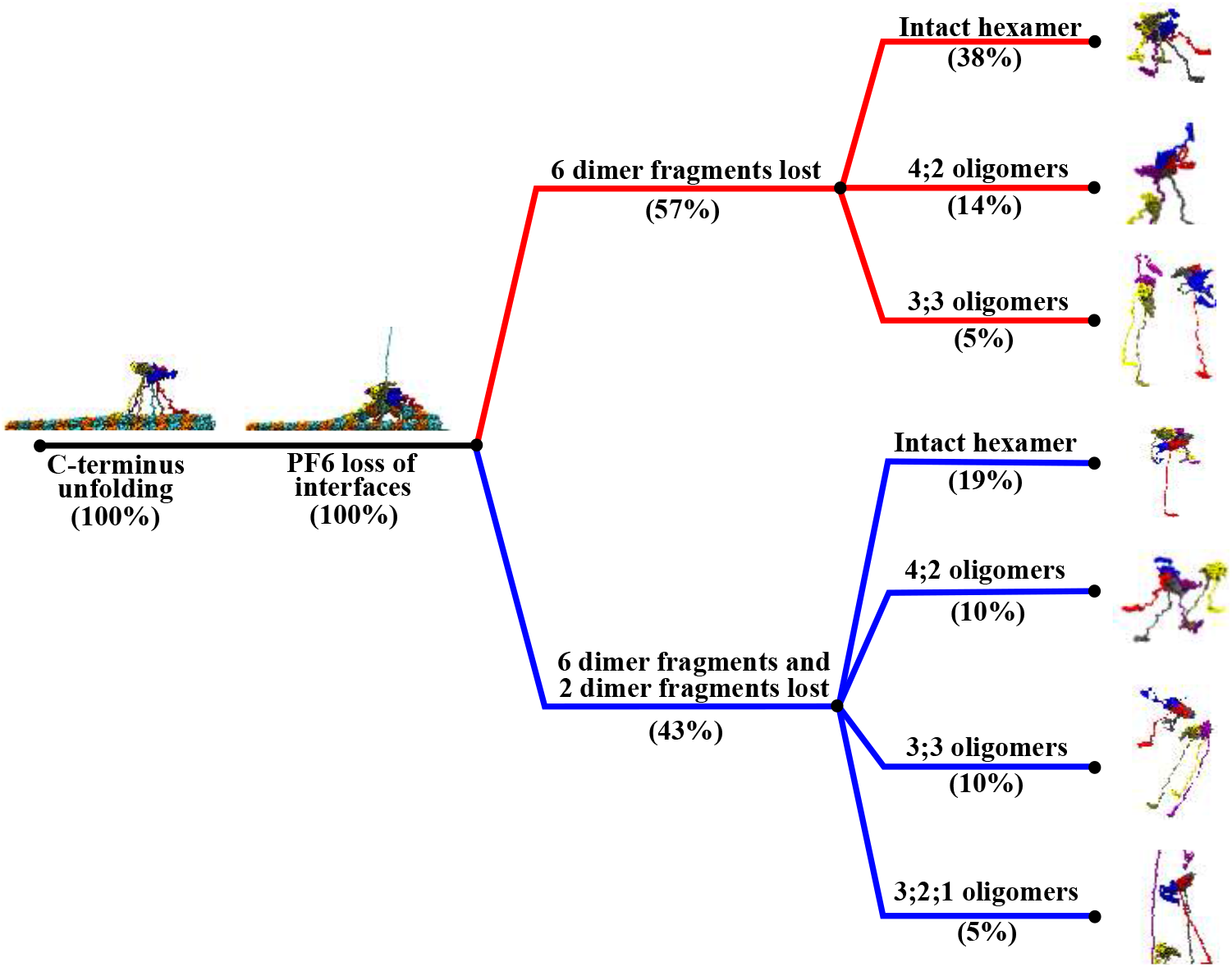
Severing pathways found for the unfoldase severing action of the full spastin machine with free MIT domains on MT8 lattice for all the probed interaction strengths between the MIT domains and the MT lattice. Descriptions and percentage of event occurrences (out of the 21 trajectories) are provided for each main event of a severing mechanism. Colors are used to separate major diverging pathways.

## 4. Direction-dependent remodeling of DHFR domains

To discern the role of force directionality in SP unfolding mediated by Clp ATPases, we comparatively probe remodeling mechanisms of three dihydrofolate reductase (DHFR) domains, one having the wild–type (WT) amino acid sequence and two with sequences obtained through circular permutation (CP), namely CP P25 and CP K38 (see Methods) [68]. Experimental approaches using wild-type and CP variants of SPs highlighted the greater importance in the unfolding process of the local SP structure near the tagged terminal over the global SP stability [69]. The identical three-dimensional structure of the SP domains, comprising four *α*-helices wrapped around a central eight-stranded *β*-sheet, affords a unique view of the effect of force directionality on unfolding mechanisms due to distinct local secondary structural elements near the terminals. The N–terminals of WT–DHFR and CP K38 consist of buried *β*–strands and the N-terminal of CP P25 consists of a solvent– exposed *α* helix, whereas the C–terminal of WT–DHFR consists of a buried *β*–strand.

We adopt an atomistic simulation model (see Methods) that probes the relative mechanical resistance of the three DHFR domains and provides detailed information on the native and non-native SP interactions involved in the unfolding and translocation processes. These processes are guided by application of stochastic forces onto the SP arising from its interaction with ClpY pore loops, which undergo repetitive axial motions during nonconcerted ATP-driven conformational transitions of ClpY subunits. To sample in a computationally efficient manner the long time scales associated with SP unfolding and translocation, our model includes application of an additional axial force onto the SP fragment located transiently within the pore region (see Methods and Table 3). The magnitude of this external force is selected such as to reflect the minimal requirement for observation of meaningful unfolding and translocation events during the computationally-accessible time scale, thereby providing a measure of the mechanical strength of each direction probed.

**Table 3.**
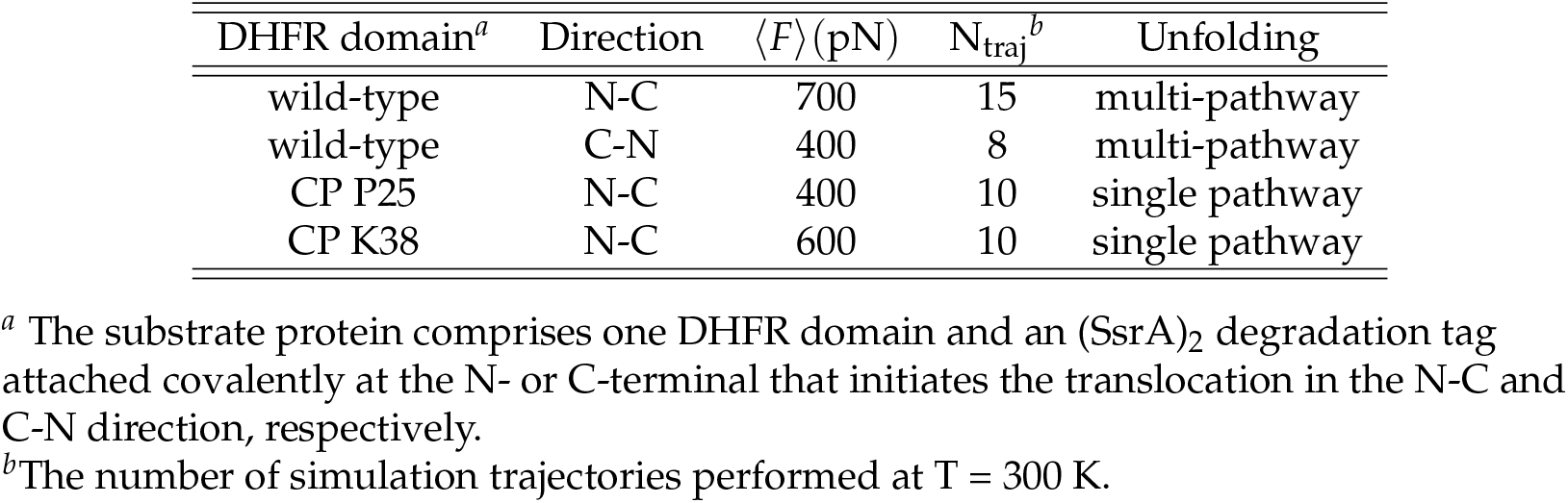
Summary of ClpYΔI-SP simulations performed.

### 4.1. Divergent unfolding pathways in WT-DHFR and CP variants

Engagement of the SP by the Clp ATPase at a single terminal and the ability of the SP to reorient at the surface of the machine result in distinct mechanical directions being probed upon pulling from the N- or C-terminals to effect N-C and C-N translocation, respectively. Consistently, we find that unfolding of the WT-DHFR domain requires a larger force, 〈*F*〉 ≃ 700 pN, when pulling is applied at the N-terminal compared with the C-terminal, 〈*F*〉 ≃ 400 pN, (Figure 5A-D). In both cases, removal of the N- or C-terminal strand from the *β*-sheet encounters strong mechanical resistance, which results in partitioning of the unfolding and translocation events into higher and lower energy barrier pathways. In the set of simulations performed in our study, the lower energy barrier pathway in N-C translocation comprises approximately 60% of trajectories that yield nearly complete unfolding, 〈Q_N_〉 ⩽ 0.3, within the simulated time frame of 400*τ*. In simulations of C-terminal pulling, the lower energy pathway comprises 50% of trajectories that yield nearly complete unfolding when the weaker pulling force is applied. Divergent mechanical resistance of the DHFR domain in N-C and C-N translocation can be understood by examining the requisite unfolding events to disrupt the tertiary structure of the domain. In N-terminal pulling, overwhelming mechanical resistance is encountered in removal of the *β*1 strand given the required disruption of the core of the *β*-sheet and of the entire domain. This major unfolding step ensnares a large fraction of the domain residues and therefore involves a drastic rewiring of the protein. This rewiring is achieved through formation of a large number of non-native contacts, 〈f_NN_〉 ≃ 0.35 (Fig. 5B), with the largest values consistent with the unfolding hindrance along the higher energy barrier pathway. In C-terminal pulling, weaker mechanical resistance is encountered in the removal of the terminal *β*10 strand as this is located closer to the protein surface. Formation of fewer non-native contacts, 〈f_NN_〉 ≃ 0.2 (Fig. 5D), allows the unfolding process to proceed with a relatively smaller pulling force requirement compared with the N-terminal case. As in the N-terminal pulling, above-average values of f_NN_ are associated with hindered unfolding and the higher energy barrier pathway.

**Figure 5.**
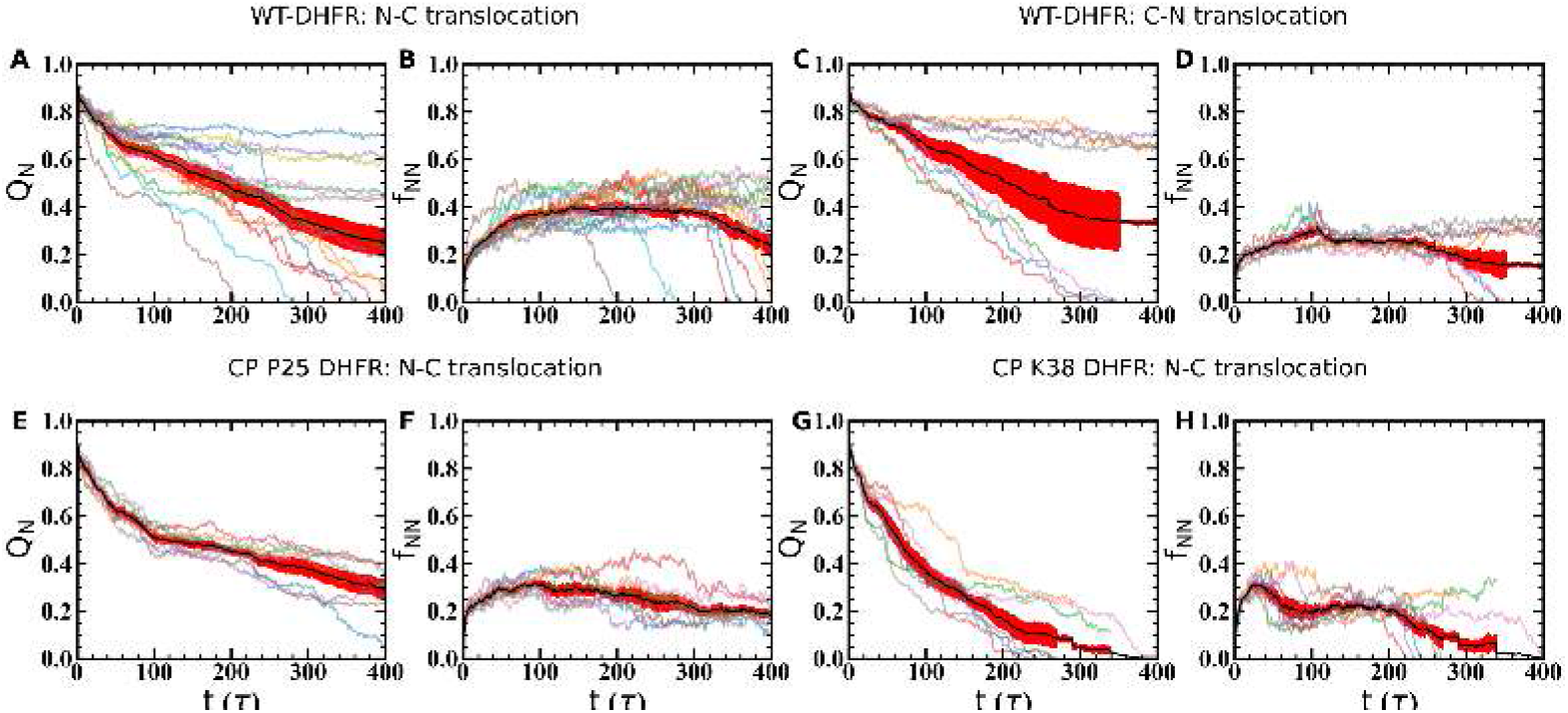
Unfolding of DHFR variants mediated by ClpY. The time evolution of the fraction of native, Q_N_, and non-native, fNN, contacts is shown for wild-type DHFR in (A)-(B) N-C and (C)-(D) C-N translocation; (E)-(F) for CP P25 and (G)-(H) for CP K38 in N-C translocation. Individual trajectories are indicated by using thin curves. Averages (thick curves, black) and standard errors (red) are also indicated.

Unfolding of the mechanical interfaces associated with the CP variants illustrate distinctive mechanisms compared with the wild-type. Pulling of the *α*-helix at the N-terminal of the CP P25 variant probes a weaker mechanical interface, therefore the application of the force of 〈*F*〉 ≃ 400 pN results in significant loss of native structure (*Q_N_* ≲ 0.5) within *t* ~ 100*τ* and unfolding along a single pathway (Fig. 5E). Interestingly, in this case, completion of the unfolding and translocation process encounters stronger mechanical resistance from the internal *β*-sheet structure and the formation of stable non-native contacts than from the N-terminal structure (Fig. 5F). Pulling at the N-terminal of the CP K38 variant requires a larger force, 〈*F*〉 ≃ 600 pN, consistent with the stronger resistance of the *β*-strand, however weaker resistance is associated with the location of the strand near the protein surface (Fig. 5G). Once the engineered N-terminal is removed from the domain core, the DHFR domain unravels rapidly, along a single pathway, without significant hindrance from non-native contacts (Fig. 5H).

### 4.2. Dynamic substrate orientation on the ATPase surface modulates the energy barrier and mechanism of unfolding

Clp nanomachine’s dynamic pore configuration and surface heterogeneity allow it to reorient the SP such that pulling is applied along favorable directions with weaker mechanical resistance. The ability to probe a variety of mechanical directions of SPs represents a tremendous advantage, especially for highly anisotropic structural elements. For instance, unfolding of a *β*-sheet may involve stronger mechanical resistance and a higher energy barrier when the pulling force is applied in the direction parallel to the strand registry, and hydrogen bonds are removed cooperatively through a shearing mechanism. By contrast, weaker mechanical resistance and a lower energy barrier for unfolding occur when the perpendicular direction is probed, and therefore inter-strand hydrogen bonds are removed sequentially through an unzipping mechanism. As shown in Figures 6, 6, and S12A-B, these aspects of mechanical anisotropy and force directionality combine to yield branched pathways in both N- and C-terminal pulling. Mechanical resistance to unfolding in the higher energy pathways is associated with larger values of the polar angle *θ* of the DHFR domain prior to translocation (translocated fraction *x* ⩽ 0.1), with significant sampling of the ranges 60 − 90°in N-terminal pulling (Figures 6A-C and 7A) and 50 − 75°in C-terminal pulling (Figures 6I-K and 7C). In the lower energy barrier pathways, orientations corresponding to 〈*θ*〉 ≃ 50°, corresponding to large Q_N_ and low x (Figures 6E-G and M-O), enable initial unfolding via unzipping and completion of the unfolding and translocation through further reorientation of the untranslocated DHFR fragment (Figures 7B and D). In these pathways, the azimuthal angle samples broad ranges, which indicates the large rotational flexibility of the DHFR domain on the surface of the Clp ATPase (Figures 6H and P).

**Figure 6.**
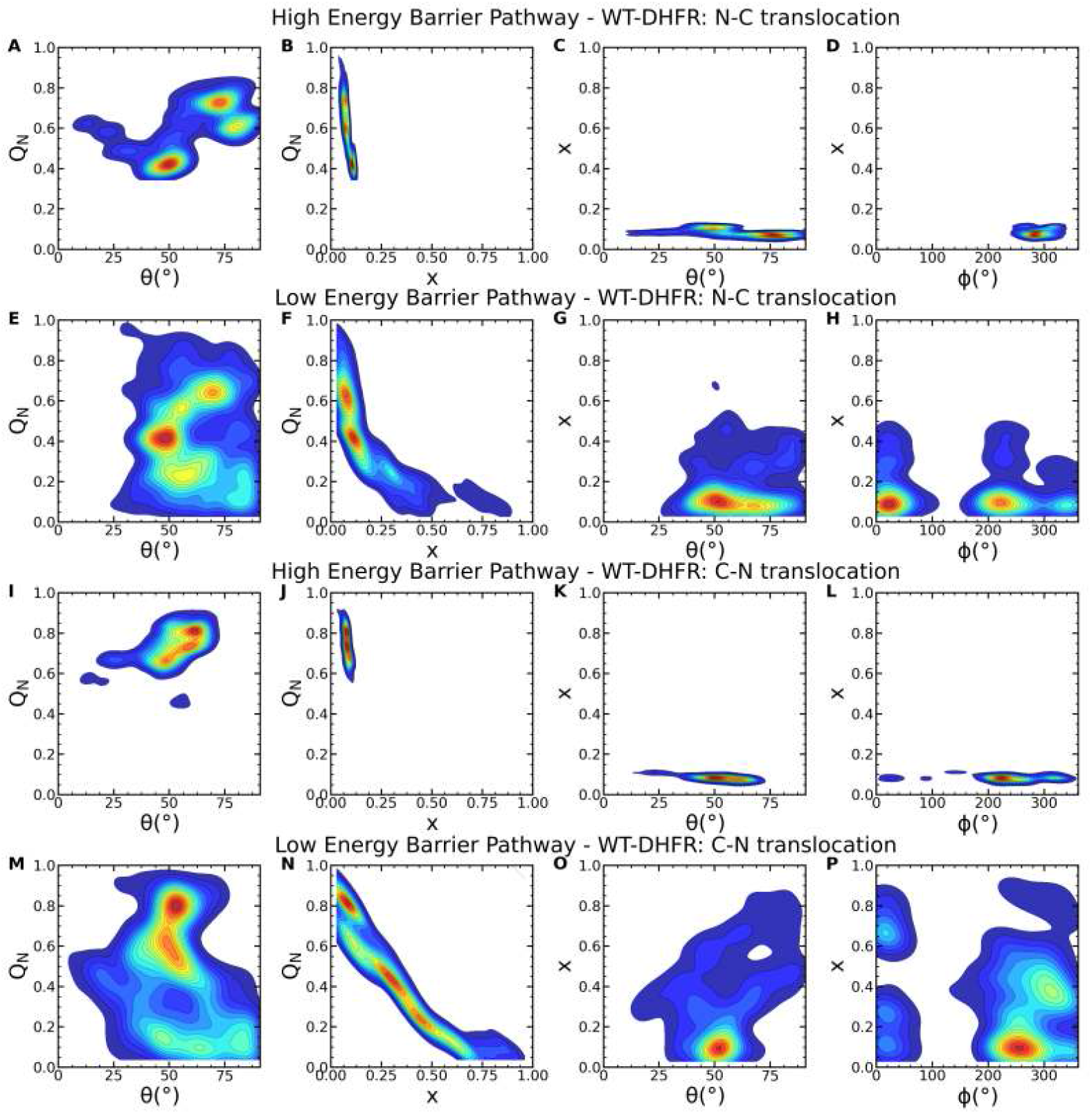
Wild-type DHFR substrate orientation at the ClpY pore lumen in unfolding and translocation pathways. Probability density maps of the (A) fraction of native contacts Q_N_ vs. polar angle *θ* and (B) translocation fraction x; (C) translocation fraction vs. polar and azimuthal (*ϕ*) angles in N-C translocation in the high-energy barrier pathway. (E)-(H) Same as in (A)-(D) in the low energy barrier pathway. (I)-(P) Same as in (A)-(H) in C-N translocation.

**Figure 7.**
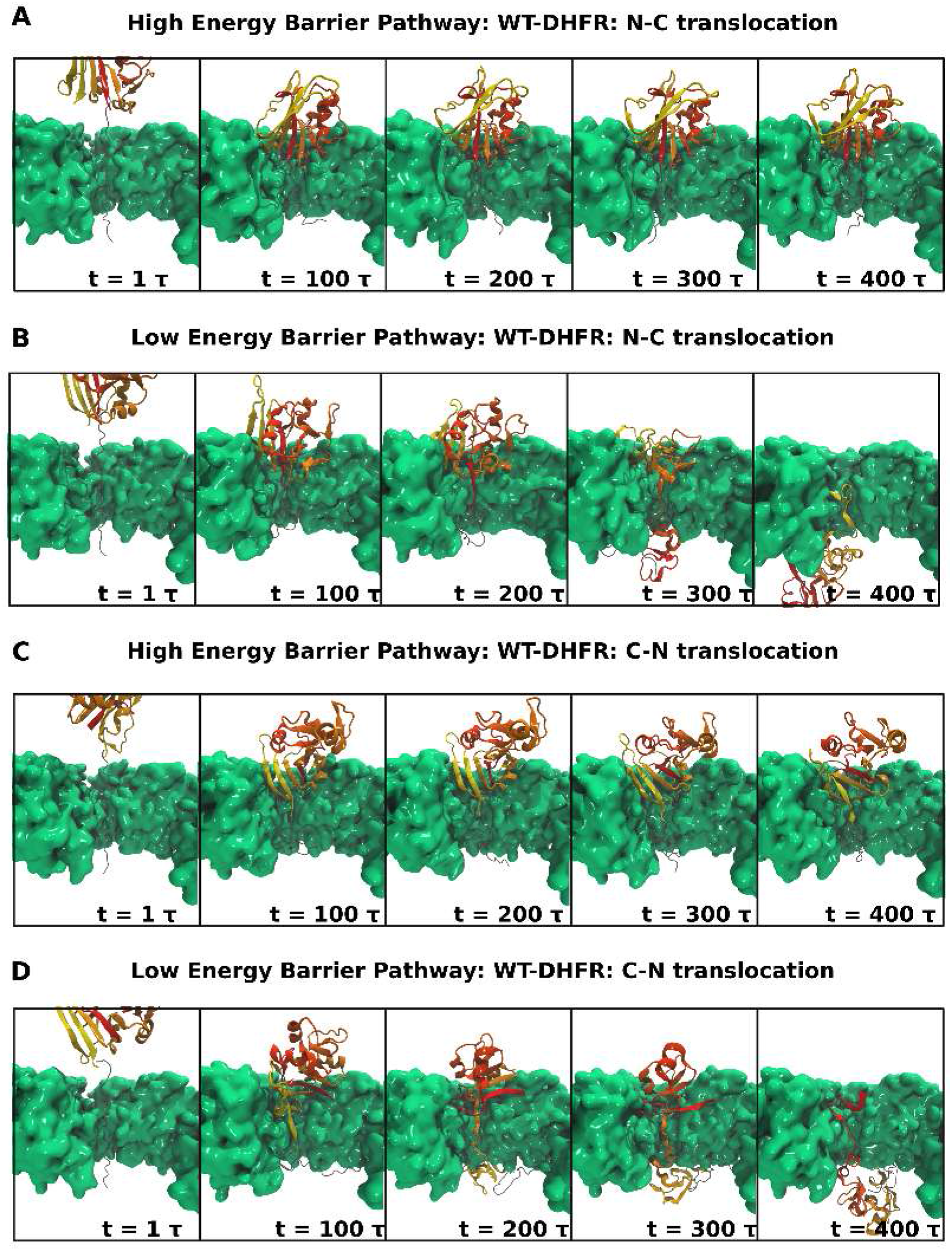
Dynamic orientation of wild-type DHFR at the ClpY pore lumen. The time-dependent orientation of DHFR (color-coded according to secondary structure) near the ClpYΔI (green) pore lumen is shown in translocation in the (A)-(B) N-C direction in the (A) high and (B) low energy barrier pathways and (C)-(D) C-N direction. Two ClpY protomers are not shown for clarity.

Unfolding of DHFR CP P25 and K38 variants involves weaker mechanical resistance, therefore their rotational flexibility has aspects that are similar to those of the low energy barrier pathways of the wild-type domain. As shown in Figures S12C and S13A-D, initial unfolding and translocation of CP P25, corresponding to large Q_N_ and low translocation fraction x, the polar angle *θ* samples the region around 50°, which correspond to orientations of the substrate such that unfolding of the *β*-sheet can be effected through an unzipping mechanism. By contrast, initial unfolding of K38 requires removal of the terminal *α*-helix, which imposes only limited restrictions to the polar angle and allows sampling of a broad range between ≃ 25 − 90°(Figures S12D and S13E-H). Both variants include large rotational flexibility indicated by the ranges of angles *θ* and *ϕ* as unfolding and translocation proceed.

### 4.3. Formation of non-native contacts modulates translocation compliance of the substrate protein

Comparative studies of the unfolding and translocation of WT-DHFR and CP variant domains, which share a common fold, allow us to examine the effect of the order of unfolding events on remodeling mechanisms. To this end, we probe the dynamic evolution of native and non-native contacts involving each secondary structural element and the characteristic dwell times associated with the unfolding steps.

As shown in Figure 8, in both N-C and C-N translocation, the evolution of native and non-native contacts is strikingly different in high- and low-energy barrier pathways. In the N-C direction, in the high-energy barrier pathway, native contacts associated with the *β*-sheet are largely preserved even as a large number of non-native contacts are formed that primarily involve *α* helices (Figure 8A-B). The strong stability of the *β*-sheet confers mechanical resistance and yields a low translocation fraction. An altogether different behavior is noted in the low-energy barrier pathway (Figure 8C-D), which reveals a substantial loss of native contacts within a time scale of ≳ 150*τ* concomitant with the dissolution of the *β*-sheet structure. Transient formation of non-native contacts is observed as the pulled polypeptide segment slides against the untranslocated DHFR fragment, but no resistance points persist over the time scale probed in our simulations. Similar behavior is found in C-N translocation, however with an even stronger divergence between the two pathways (Figure 8E-H).

**Figure 8.**
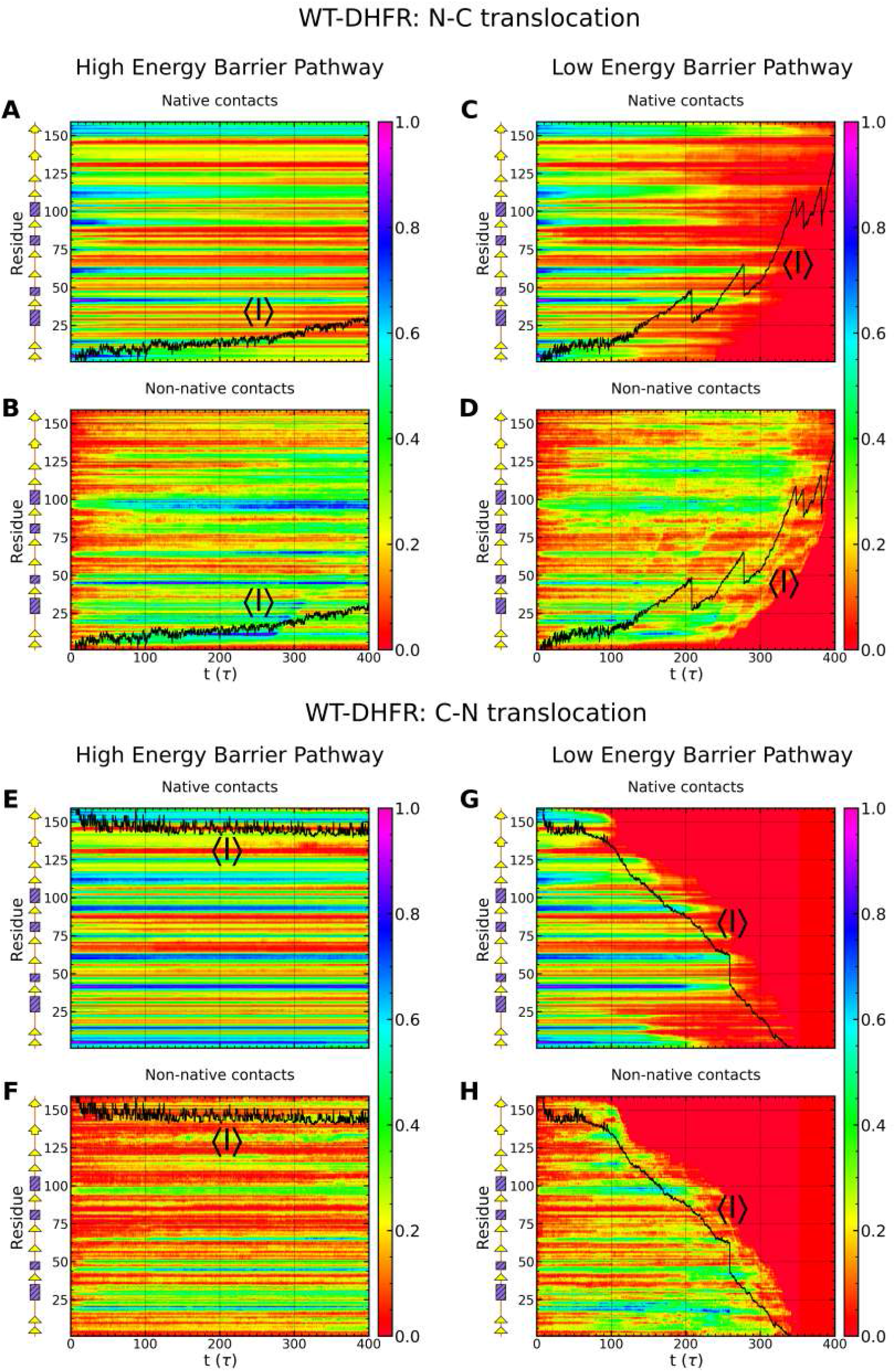
Direction-dependent time evolution of native and non-native content of wild-type DHFR. (A)-(D) The time-dependent fraction of native Q_N_ and non-native fNN contacts formed by secondary structural elements of wild-type DHFR during translocation in the N-C direction in the (A)-(B) high and (C)-(D) low energy barrier pathway. The time–dependent average translocation line (black) is also indicated.

Native and non-native contacts of CP variants are more compliant than those of WT-DHFR in the initiation of Clp-mediated unfolding. As shown in Figures 9A-B, native contacts near the engineered N-terminals are lost on a time scale of 100 *τ* and transient non-native contacts in this region are formed primarily within the P25 variant. Nevertheless, as noted above, complete unfolding and translocation of the P25 variant encounters strong mechanical resistance once the core *β*-sheet (*β*1 − *β*2 and *β*7 − *β*10 strands) is directly engaged. Native contacts in this region persist over long times with limited contribution of non-native contacts.

**Figure 9.**
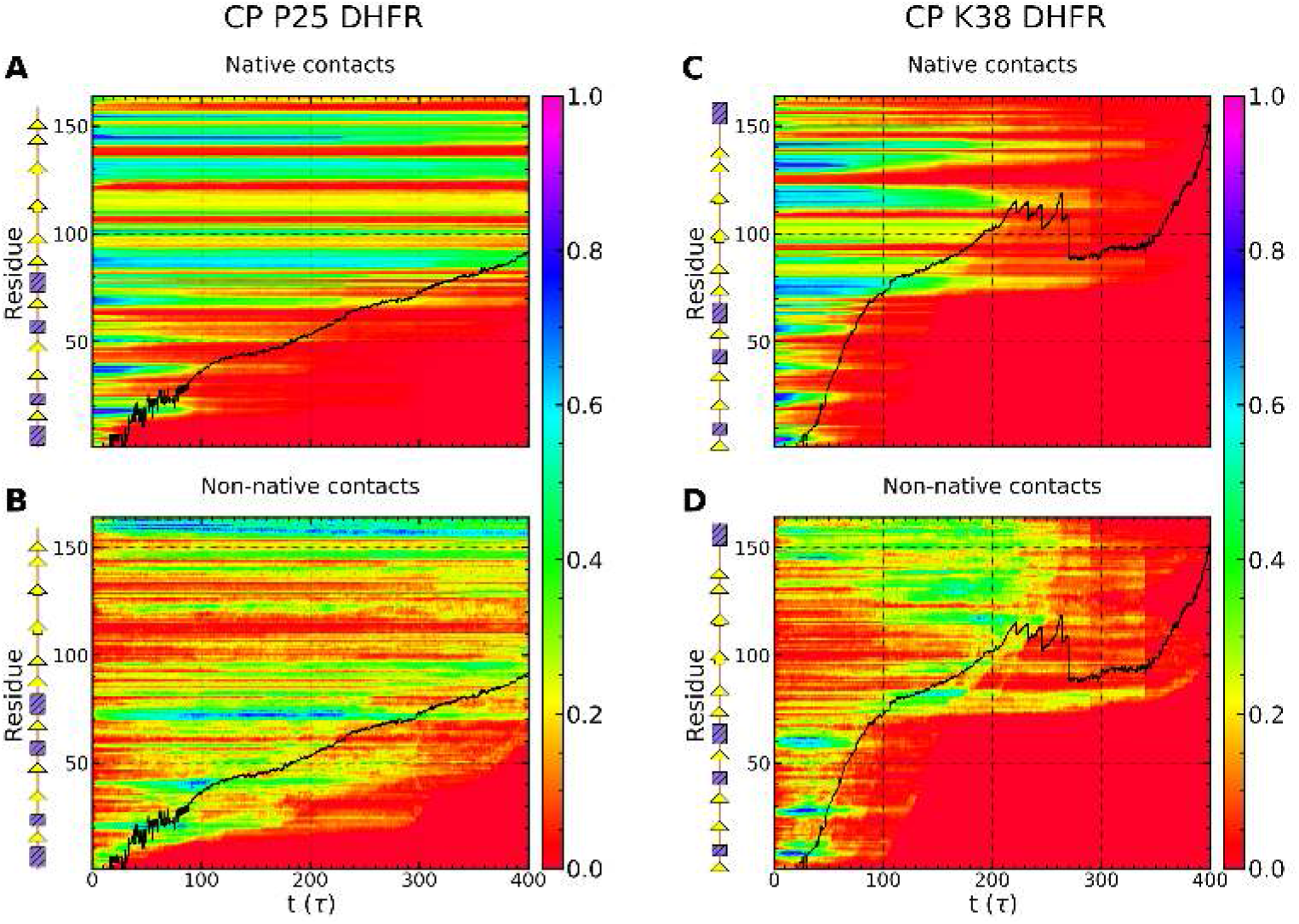
Direction-dependent time evolution of native and non-native content of circular permutant variants P25 and K38 of DHFR. Representations are the same as in Figure 8.

Diverging dynamic contribution of native and non-native contacts of the WT-DHFR and CP variants highlights the importance of kinetic aspects of the unfolding and translocation processes. Quantitatively, these kinetic aspects can be characterized by determining the waiting time per residue during the translocation process that provides a fingerprint of the mechanical resistance of the polypeptide chain (see Methods) [34,70,71]. As shown in Figure 10, in the high-energy barrier pathways, in both N- and C-terminal pulling of the WT-DHFR, large dwell times reflect the strong mechanical resistance near the engaged terminal. These large barriers are so large that they cannot be overcome during the time scales of our simulations, therefore no significant translocation is observed in these pathways. In the low-energy barrier pathways, smaller barriers are associated with the initial unfolding events, which enable the Clp ATPase to overcome them through repetitive force application and facilitate complete translocation as small internal barriers are subsequently encountered. Translocation of CP variants does not involve any significant dwelling upon the initial SP engagement by the Clp ATPase, however one or more internal barriers are encountered that result in larger waiting times. These dwell times associated with these partially unfolded intermediates culminate with those found upon removal of all the SP structure except for the *β*-sheet. In the case of CP P25, this results in translocation times longer than even the time scales probed in our simulations, whereas in the case of CP K38 they yield very long times for complete unfolding and translocation.

**Figure 10.**
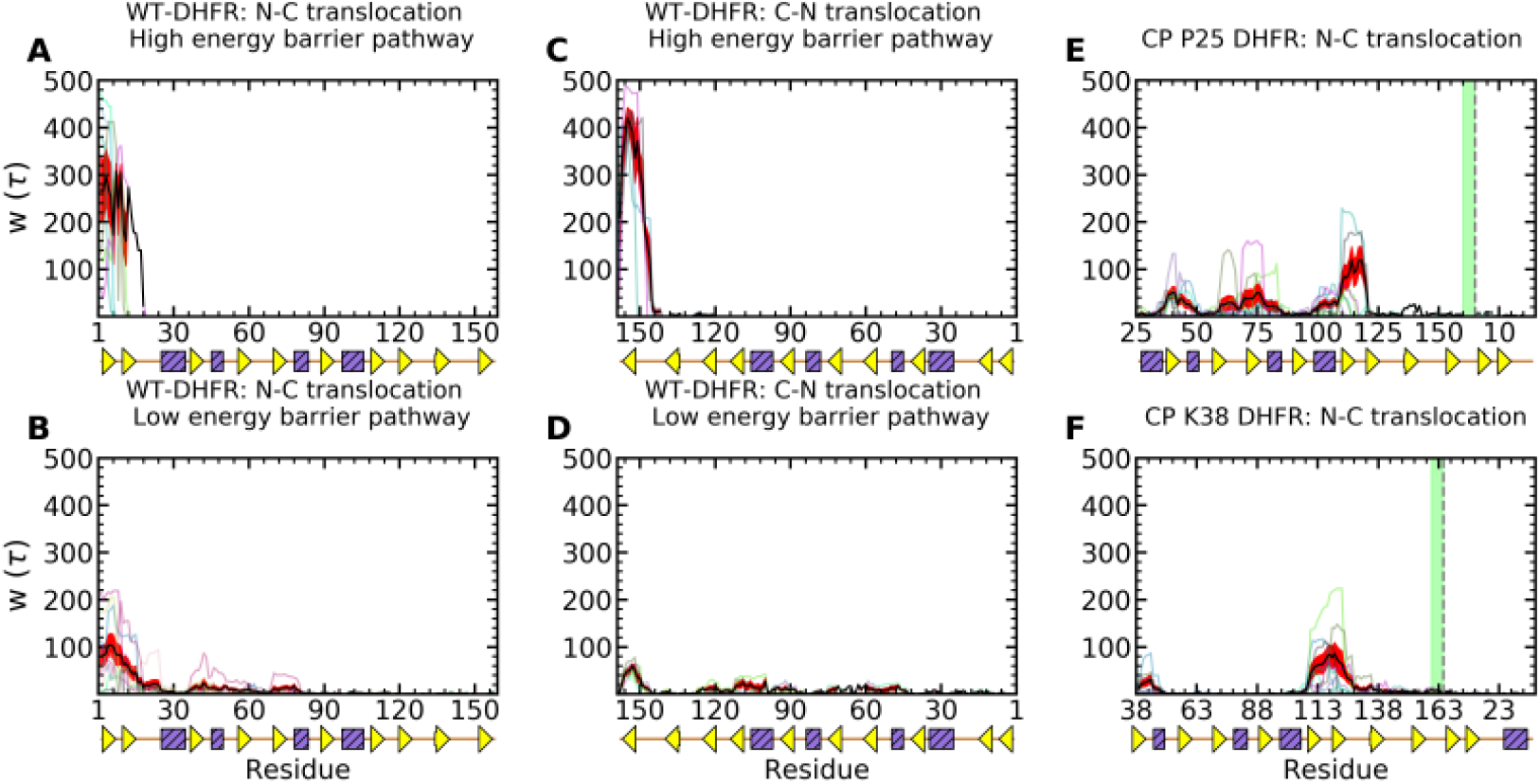
Translocation hindrance of DHFR variants. The waiting time per residue of the wild-type DHFR is shown in translocation in the (A)-(B) N-C direction in the (A) high and (B) low energy barrier pathways, in the (C)-(D) C-N direction. (E)-(F) Same as in (A) for the circular permutant DHFR variants (E) P25 and (F) K38 (the wild-type N- and C-terminal sequence positions are indicated using dashes and a green line, respectively). Traces for individual trajectories are shown using thin curves and averaged values using thick black curves Standard errors are shown using red bands.

## 5. Conclusions

In these studies, we examined comparatively the unfoldase action of two representative AAA+ machines, spastin, which is responsible for microtubule-severing, and ClpY, which mediates protein degradation. The remarkable span of length scales of SPs remodeled is in strong contrast to the structural and functional similarities of the two types of nanomachines. Both spastin and ClpY have a homohexameric AAA ring structure and pulling of the substrate is effected through ATP-driven motions of central pore loops. The dramatic difference in unfolding requirements of SPs found at the extremes of these broad length scales prompted us to address the question of what underlying mechanisms endow these nanomachines with the versatility to process such diverse substrates. Given the struc-tural and functional similarity of the nanomachines, a plausible mechanism, which we set to examine in this paper, is that the dynamic relative orientation of the nanomachine and SP allows the application of mechanical force along favorable directions of weak mechanical resistance.

To effectively probe the mechanisms of the two nanomachines at the appropriate length scales, we performed molecular dynamics simulations using coarse-grained and atomistic descriptions, respectively, that allowed efficient sampling of the conformational space in each case. To this end, we developed coarse-grained models of the spastin-microtubule system that included either the truncated spastin, comprising the motor domain, or the complete machine, comprising the MIT domain, the motor domain and the linkers. Comparison between the unfolding requirements in the two spastin setups provide a detailed understanding of the contribution of the motor and MIT domains to the unfoldase function. We also developed atomistic models of the ClpY-SP system that probed the direction-dependent unfolding of DHFR by considering wild-type and CP variants. These simulations highlight the dependence of SP unfolding and translocation pathways on both the local mechanical strength near the engaged terminal and the internal wiring of the substrate. Overall, the emerging conclusion is that complete unfolding and translocation requires unhindered ability to apply force along softer mechanical directions in order to overcome both the native contacts that stabilize the mechanical interfaces that experience mechanical pulling and the non-native contacts that dynamically form along each pathway. Our studies of the breaking of a MT lattice by severing enzymes showed variations in the unfoldase mechanism based on the types of parameters set for our model [43]. Initial pulling simulations involving the fixing of N-terminal residues in the spastin motor mounted on a MT lattice resulted in the removal of the pulled monomer, after its substantial unfolding, when the motor was kept anchored on the MT surface or of the pulled tubulin dimer when only selected spastin monomers were fixed on the lattice. The latter finding suggests a potential functional role for the linkers and MIT domains of allowing more room for the orientation of the spastin motor with respect to the MT lattice. To test this proposal, we modeled the full spastin machine, including the MIT domains attached to the ATPase motor through flexible linkers, such that the motor would now be free to reorient on the MT filament. Overall, these runs resulted in free fluctuations of the MIT domains and MT breaking patterns reminiscent of our earlier findings for the pulling on a single dimer with the MT lattice plus end free [43]. Varying the interaction strength between the severing machine and the MT lattice showcased two characteristics of the spastin machine found in the literature. For a weak interaction strength (1.0 and 1.5 kcal/mol) [23], the machine detaches intact with the pulled fragment from the MT filament. By contrast, at interaction strengths 2.5 kcal/mol, which correspond to the binding affinity of kinesin-1 motors on MTs [67], we found that, after breaking multi-dimer PF fragments, the spastin machine usually dissociates into 2 trimers, while remaining in contact with the MT filament. For even stronger interactions, the machine usually dissociates into smaller oligomers, which remain on the MT surface upon the loss of multi-dimer PF fragments. This behavior was found only at interaction values above 2.00 kcal/mol, and is of interest due to the proposal from the literature that spastin and other members from the same AAA+ clade (katanin and ClpB) undergo dissociation upon removal of the substrate and the ATP [72,73]. These findings are important in understanding the oligomerization and unfolding action of severing proteins. Regarding the orientation of the severing machine versus its substrate, our simulations showed that lower interaction strengths lead to large movements of the spastin machine, and the alignment of the central pore loop’s along the axis of the pulled PF. As the strength between the MIT domains and the lattice increased, the spastin motor and the full machine maintained their orientation with respect to the MT filament for most of the run until severing of a PF fragment is achieved. The critical breaking force leading to the cutting of PF fragment reached its highest value when the motor could orient parallel with the MT, which occurred at lower interaction strengths. These observations, obtained by modeling GDP microtubule lattices with interactions between subunits at the level of 1.0 kcal/mol, have shown overall trends found in the severing mechanisms. It is important to note that the major factor which dictates the severing pathways is the relative strength of the interactions between the spastin machine and the MT filament versus the strength of the intra- and inter-protofilament contacts of the MT. An important aspect regarding the MT cytoskeleton is the tubulin code [74], which refers to the fact that, while all MTs are polymeric assemblies of tubulin dimers, there are many post-translational modifications on the tubulin monomers as well as tubulin isotypes which control the properties and functions of MTs. For example, acetylation, a post-translational modification that occurs in tubulin, alters the rigidity of the lattice compared to the standard GDP lattice by decreasing the interaction strength between protofilaments and would therefore modify the *ϵ_h_* between the MIT domains of spastin and the lattice [74]. Similar changes can be envisioned due to changes in the tubulin sequence based on the cell type or organism [75]. This, in turn, would lead to altered propensity of the breaking pathways. In Clp-mediated remodeling, the counterpart action to changes in the orientation of the severing machine versus the MT filaments resulting from differences in the rigidity of the filaments due to the tubulin code consists in the selection by the machine of specific orientations relative to each DHFR variant. The fate of the globular protein upon engagement by the Clp ATPase is largely determined by the mechanical resistance offered by the local structure. Nevertheless, while the mechanical strength of the relevant interface can be inferred from the type of secondary structure present, with *α*-helices expected to provide softer resistance compared with *β*-sheets, or the extent of solvent exposure, with interfaces near the protein surface softer compared with buried ones, our simulations of DHFR CP variants reveal that the direction of force applied by the ATPase strongly modulates the unfolding pathways. In these variants, unfolding of the core *β*-sheet involves pathways with high energy barriers when dynamic orientation of the SP at the ClpY pore lumen restricts force application to strong mechanical directions due to the location of terminal *β*-strands within the sheet, as it is the case for the wild-type N- and C-terminals. Notably, the SP orientation plays a role not only in unfolding the native structure, but also metastable intermediate conformations encountered along the degradation pathway, as illustrated by the case of the P25 CP variant, which requires unfolding of the *β*-sheet as a downstream event. In the crowded cellular environment, force directionality can be constrained by external factors, as rotational diffusion of the SP itself may be hindered due to the presence of proteins that are not directly interacting with the nanomachine. An example of rotational hindrance, illustrated by our previous simulations of multi domain I27 substrates remodeled by Clp ATPases, is the restricted rotation of the domain engaged directly by the nanomachine as other domains crowd the pore lumen [41]. The role of non-native interactions strongly separates mechanisms of microtubule-severing and protein unfolding and translocation. We expect that, given the size of the substrate (the MT lattice) and the fact that severing is not characterized by the unfolding of the tubulin monomers, the formation of any non-native interactions is likely to be transitory and thus play little to no role in the response of the MT filament to the action of spastin motors. This also represents the basis of our coarse-grained modeling approach that does not include non-native interactions between amino acids in our studies of the unfoldase action of spastin on MT lattices. By contrast, atomistic modeling used in ClpY simulations highlighted the strong contribution of non-native interactions to increased mechanical resistance of SPs that yields pathways with high energy barriers to unfolding and translocation. In general, in the Clp-mediated unfolding of globular proteins, loss of the native structure can lead to distinct mechanisms that involve formation of non-native contacts to varied degrees. At one extreme, cooperative loss of native structure yields domain unfolding through a two-state model and results in limited formation of non-native contacts. Translocation of the unfolded chain is then controlled only by the possible formation of stabilizing contacts with auxiliary domains of the machine [76]. At the other extreme, the SP unfolding process can be severely hindered by formation of strong non-native contacts. An apt illustration of this situation is provided by the ClpY-mediated unfolding and translocation of knotted SPs for which mechanical pulling drives the sliding of the knot towards the free terminal of the polypeptide chain. Deep knots have initial boundaries far (>30 residues) from the free terminal, which renders them non-compliant with sliding off the chain upon force application. As shown in our previous simulations, when the knot slides over the polypeptide chain it encounters a rough conformational landscape presented and its progress can be completely stalled by formation of non-native contacts involving side chains. Results of our comparative studies of the direction-dependent mechanisms employed by microtubule-severing and protein degradation machines, together with the structural similarity characterizing the AAA+ superfamily, raise the intriguing possibility of engineering these nanomachines for novel functions that expand on their cellular actions. For example, can a powerful double-ring machine, such as Hsp104, be repurposed to perform microtubule-severing? The demonstrated engineering of the ClpB to perform degradation within the BAP construct makes it plausible that alternative microtubule-severing nanomachines and mechanisms can be obtained. These considerations, along with the overall observations emerging from our studies, are broadly relevant for the diverse substrates remodeled by AAA+ nanomachines. In particular, the understanding of mechanisms of disaggregation of ordered fibrillar aggregates, which lie at an intermediate length scale between globular proteins and microtubules, mediated by Hsp104, is combine both aspects of ATPase-SP orientation and mechanical anisotropy of the SP. Fibrillar aggregates are stabilized by cross-*β* interactions formed between the peptides, therefore the orientation of the machine relative to the fibrillar axis controls its ability to remove monomers using an unzipping mechanism.

## Supplementary Materials

The following are available online at https://www.mdpi.com/article/10.3390/1010000/s1, Figure S1: Force vs frame number profile for the action of the spastin hexameric motor on a MT fragment, Figures S2 to S5: Results of the spastin machine acting on a MT filament when the N-terminal ends of all the MIT domains are fixed on the MT surface, Figure S6: Force vs frame number profile for the action of the spastin hexameric machine on a 8 dimers long, 13PF MT lattice, Figures S7 to S11: Results of the spastin machine acting on a MT filament for the interaction strength between its MIT domains and the MT lattice set to 1.5 kcal/mol, 2.0 kcal/mol, 3.0 kcal/mol, 3.5 kcal/mol, and 4.0 kcal/mol, respectively, Figure S12: Clustering analysis of DHFR conformations and orientations in ClpY-mediated unfolding and translocation pathways, Figure S13: CP DHFR variant orientation at the ClpY pore lumen in unfolding and translocation pathways.

## Author Contributions

Conceptualization, G.S. and R.I.D; methodology, R.A.V., H.Y.Y.F., M.S.K., M.D., G.S., and R.I.D; data collection and analysis, R.A.V., H.Y.Y.F., M.S.K., M.D., S.M., J.L.N., C.M.G., G.S., and R.I.D; writing—original draft preparation, R.A.V., H.Y.Y.F, M.S.K., G.S. and R.I.D.; supervision, G.S. and R.I.D.; funding acquisition, G.S. and R.I.D. All authors have read and agreed to the published version of the manuscript.

## Institutional Review Board Statement

Not applicable.

## Informed Consent Statement

Not applicable.

## Data Availability Statement

Not applicable.

## Acknowledgments

We thank Sue Wickner and Mike Maurizi for stimulating discussions on Clp-mediated protein degradation.

