## Supplemental Material for "Exploring the effect of mechanical anisotropy of protein structures in the unfoldase mechanism of AAA+ molecular machines"

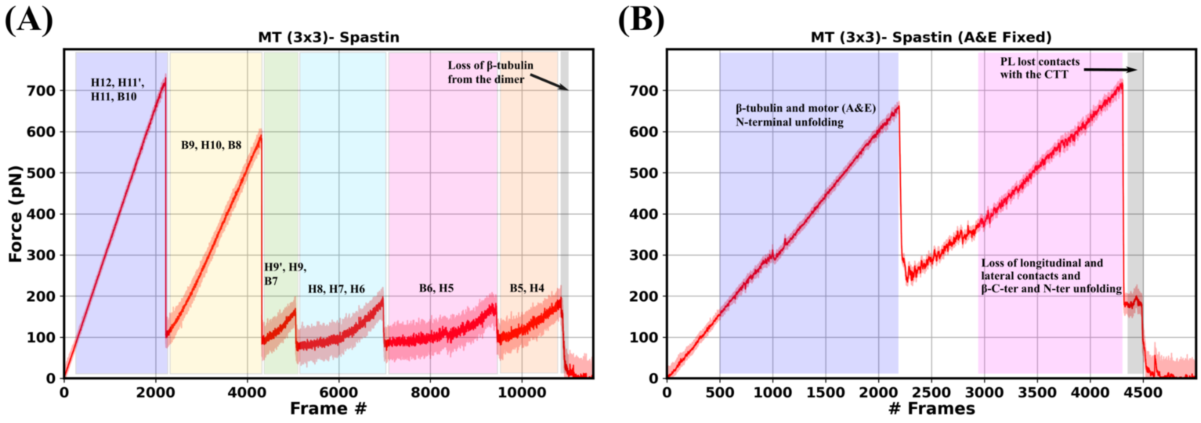


Figure S1: Force vs frame number profile for the action of the spastin hexameric motor on a $MT_{3x3}$ lattice. Here, 1 frame = 0.04 ms. The motor acts by pulling on the C-terminal end of the central β monomer when the N-term residues of (A) all spastin chains are fixed, and of (B) only spastin chains A and E are fixed.

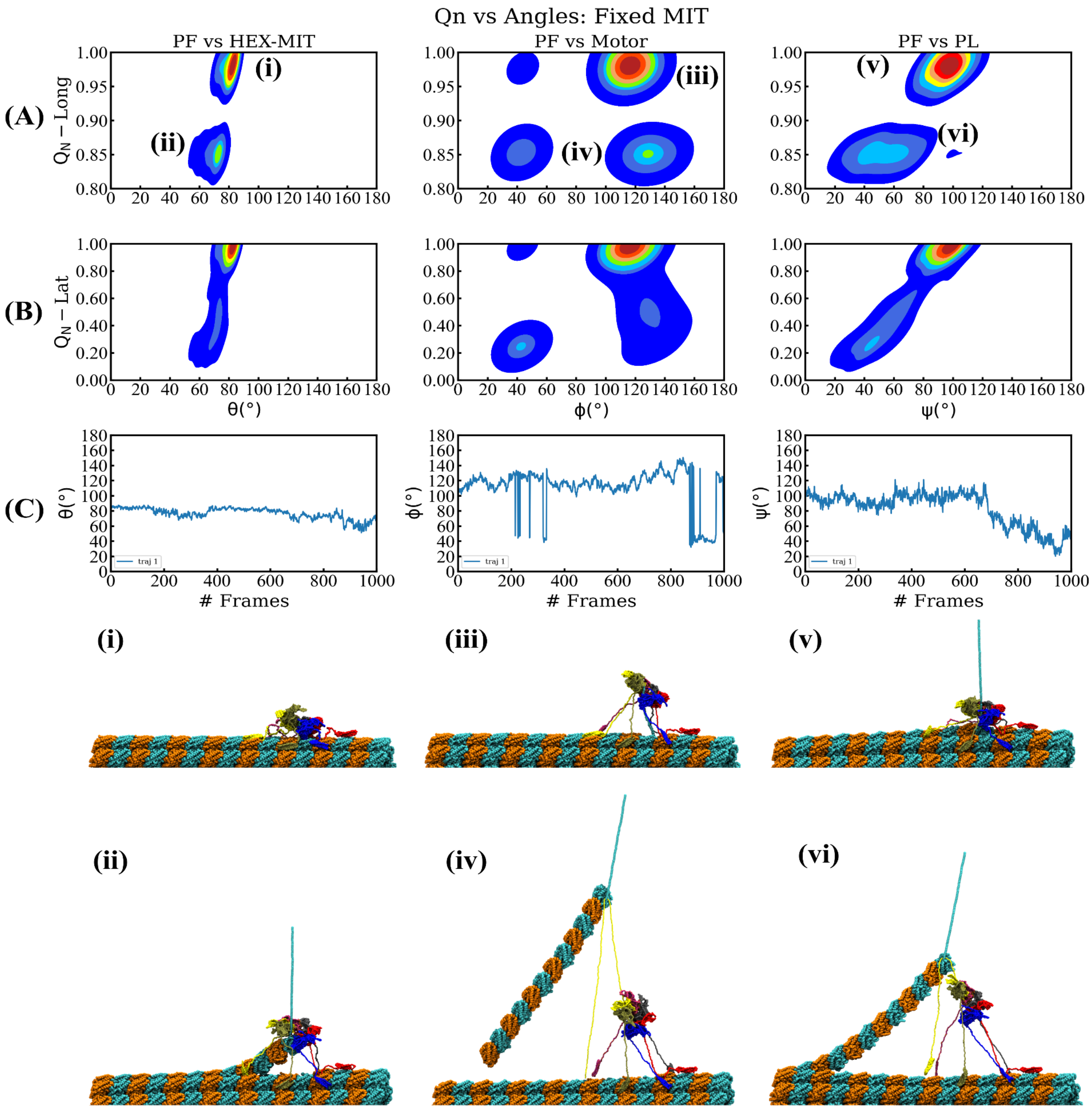

Qn vs Angles: Fixed MIT
PF vs HEX-MIT
PF vs Motor
PF vs PL
(A)
(B)
(C)
$Q_N$ − Long
$Q_N$ − Lat
θ(°)
ϕ(°)
ψ(°)
### Frames
traj 1
(i)
(ii)
(iii)
(iv)
(v)
(vi)

Figure S2: Results of the spastin machine acting on a MT filament when the N-terminal ends of all the MIT domains are fixed on the surface of the MT lattice and the interaction between the spastin machine and the MT lattice is 1.0 kcal/mol. This is similar to Fig. 2 in the main text.

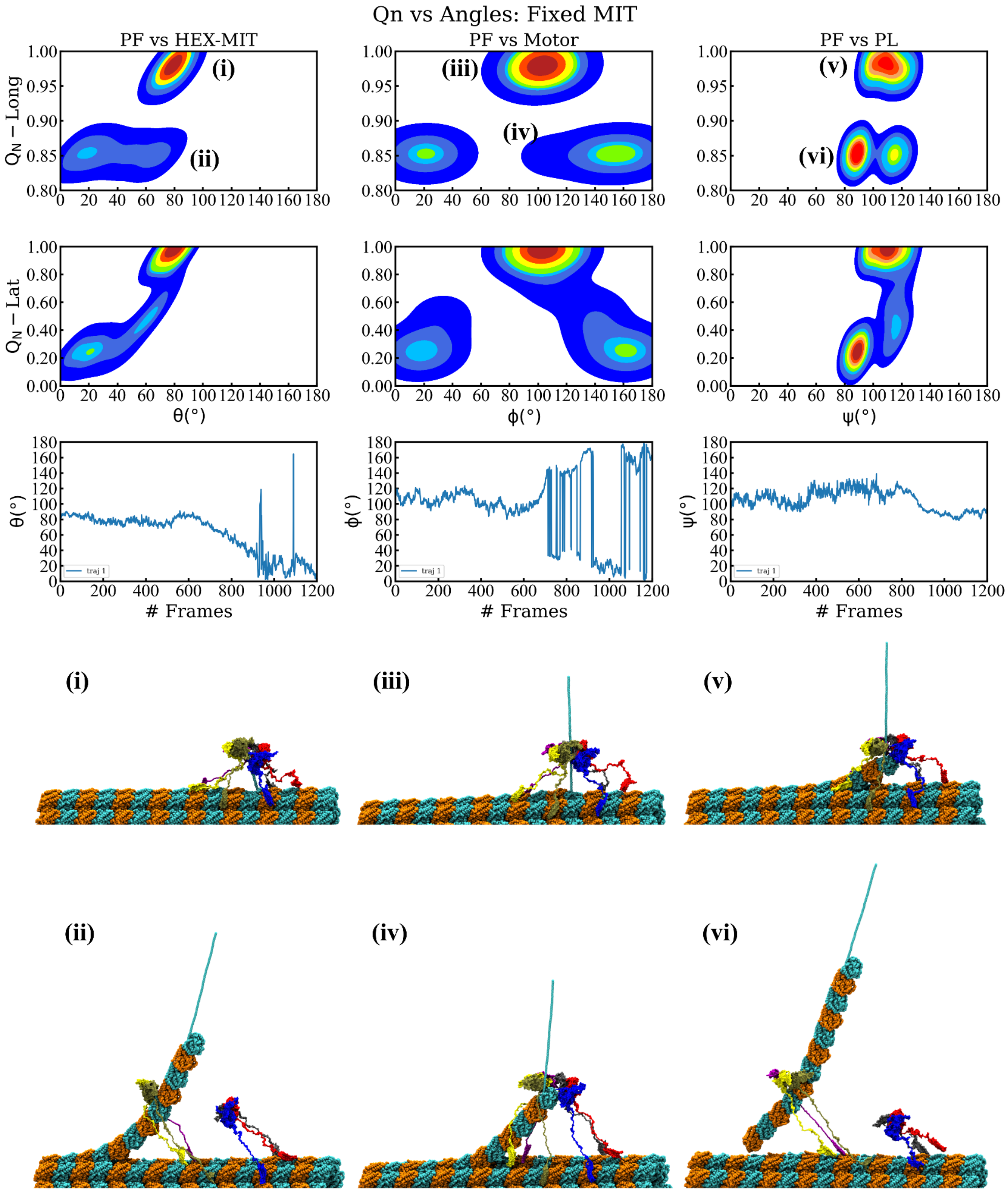

Qn vs Angles: Fixed MIT
PF vs HEX-MIT
PF vs Motor
PF vs PL
Q_N − Long
Q_N − Lat
θ(°)
ϕ(°)
ψ(°)
### Frames
traj 1
(i)
(ii)
(iii)
(iv)
(v)
(vi)

Figure S3: Similar to Fig. S2, while only the contacts between the E15 peptide at the C-terminal end of the pulled tubulin monomer and spastin are set to 1.0 kcal/mol.

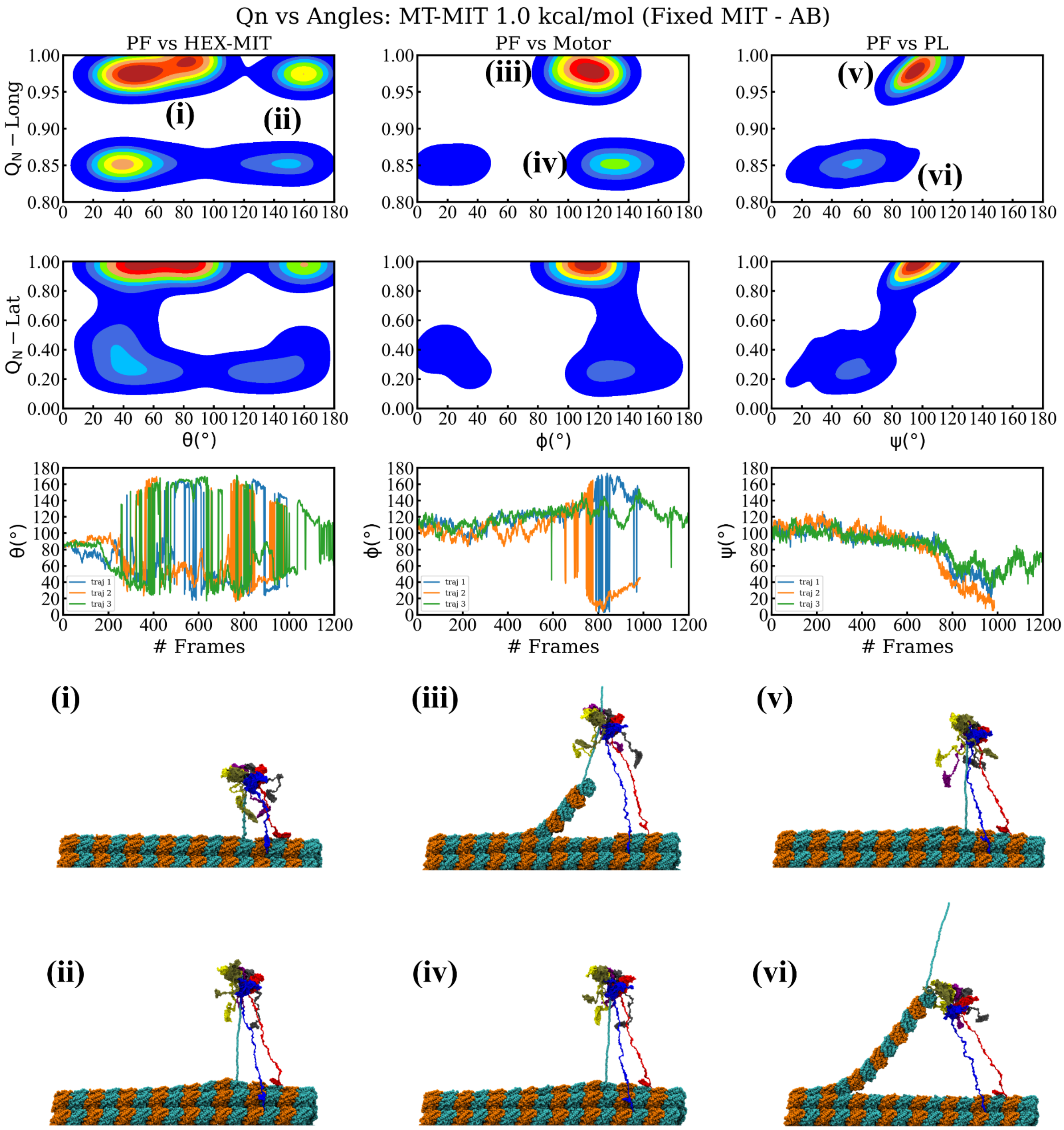
Qn vs Angles: MT-MIT 1.0 kcal/mol (Fixed MIT - AB)
PF vs HEX-MIT
PF vs Motor
PF vs PL
$Q_N$ − Long
$Q_N$ − Lat
θ(°)
ϕ(°)
ψ(°)
### Frames
traj 1
traj 2
traj 3
(i)
(ii)
(iii)
(iv)
(v)
(vi)

Figure S4: Similar to Fig. S2, but when fixing only the N-terminal ends of the MIT domains from chains A and B in spastin.

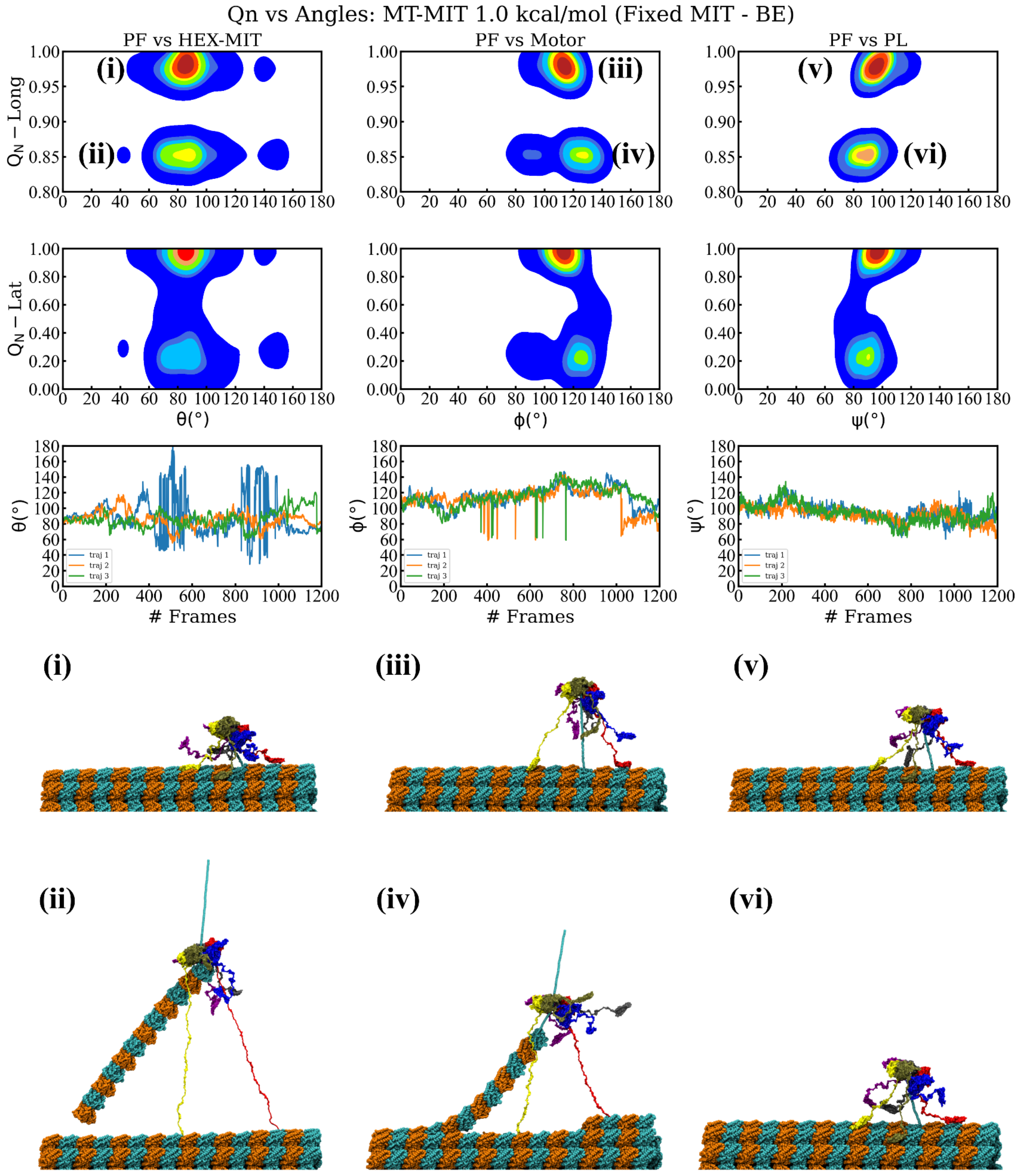

Qn vs Angles: MT-MIT 1.0 kcal/mol (Fixed MIT - BE)
PF vs HEX-MIT
PF vs Motor
PF vs PL
$Q_N$ – Long
$Q_N$ – Lat
θ(°)
ϕ(°)
ψ(°)
### Frames
traj 1
traj 2
traj 3
(i)
(ii)
(iii)
(iv)
(v)
(vi)

Figure S5: Similar to Fig. S2, but when fixing only the N-terminal ends of the MIT domains from chains B and E in spastin.

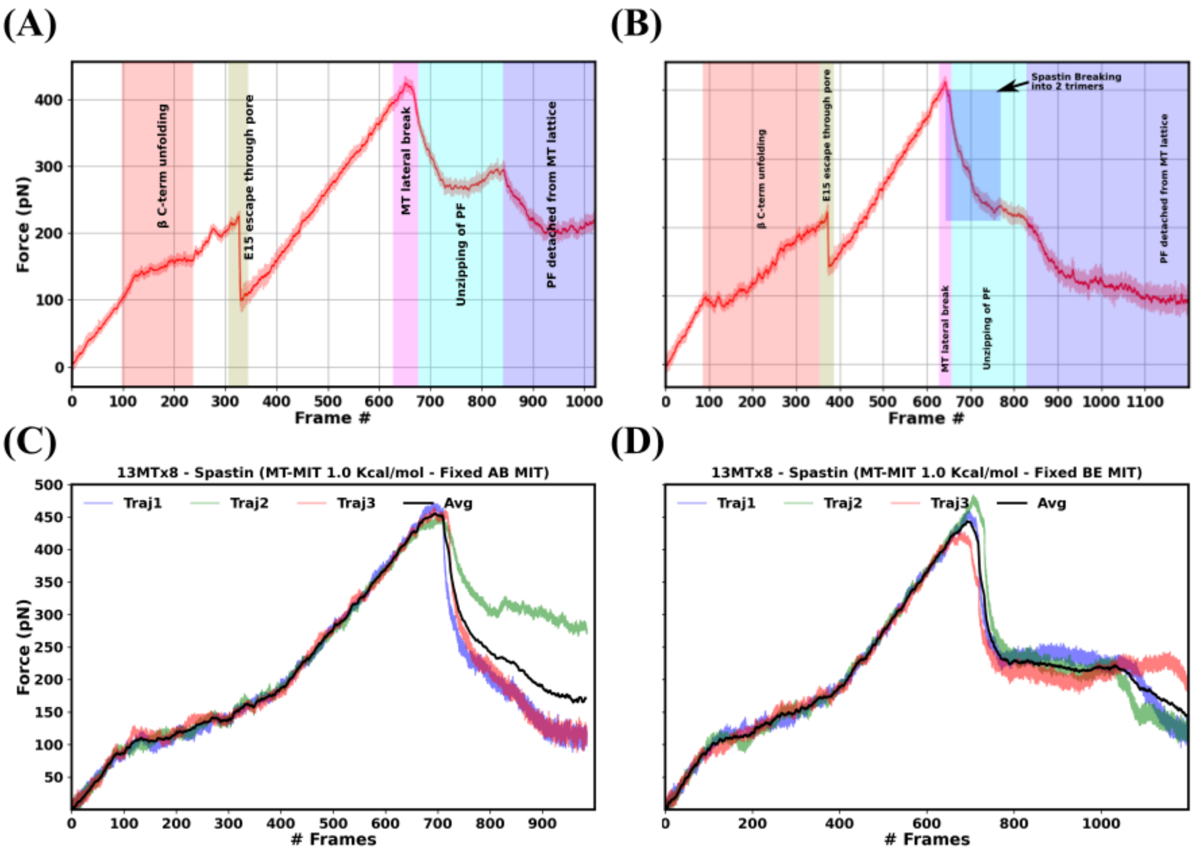


Figure S6: Force vs frame number profile for the action of the spastin hexameric machine on a 8 dimers long, 13PF MT lattice when keeping the N-terminal ends (A) of all MIT domains fixed on the MT lattice and all contacts between the MT and HEX-MIT are set to 1.0 kcal/mol, (B) of all MIT domains fixed on the MT lattice and only the contacts between the MT and MIT, and between the MT and PL are set to 1.0 kcal/mol, (C ) of only the MIT domains from two consecutive protomers (chains A and B) fixed and only the contacts between the MT and MIT, and between the MT and PL are set to 1.0 kcal/mol, and (D) of only the MIT domains from two opposite protomers (chains B and E) fixed and only the contacts between the MT and MIT, and between the MT and PL are set to 1.0 kcal/mol.

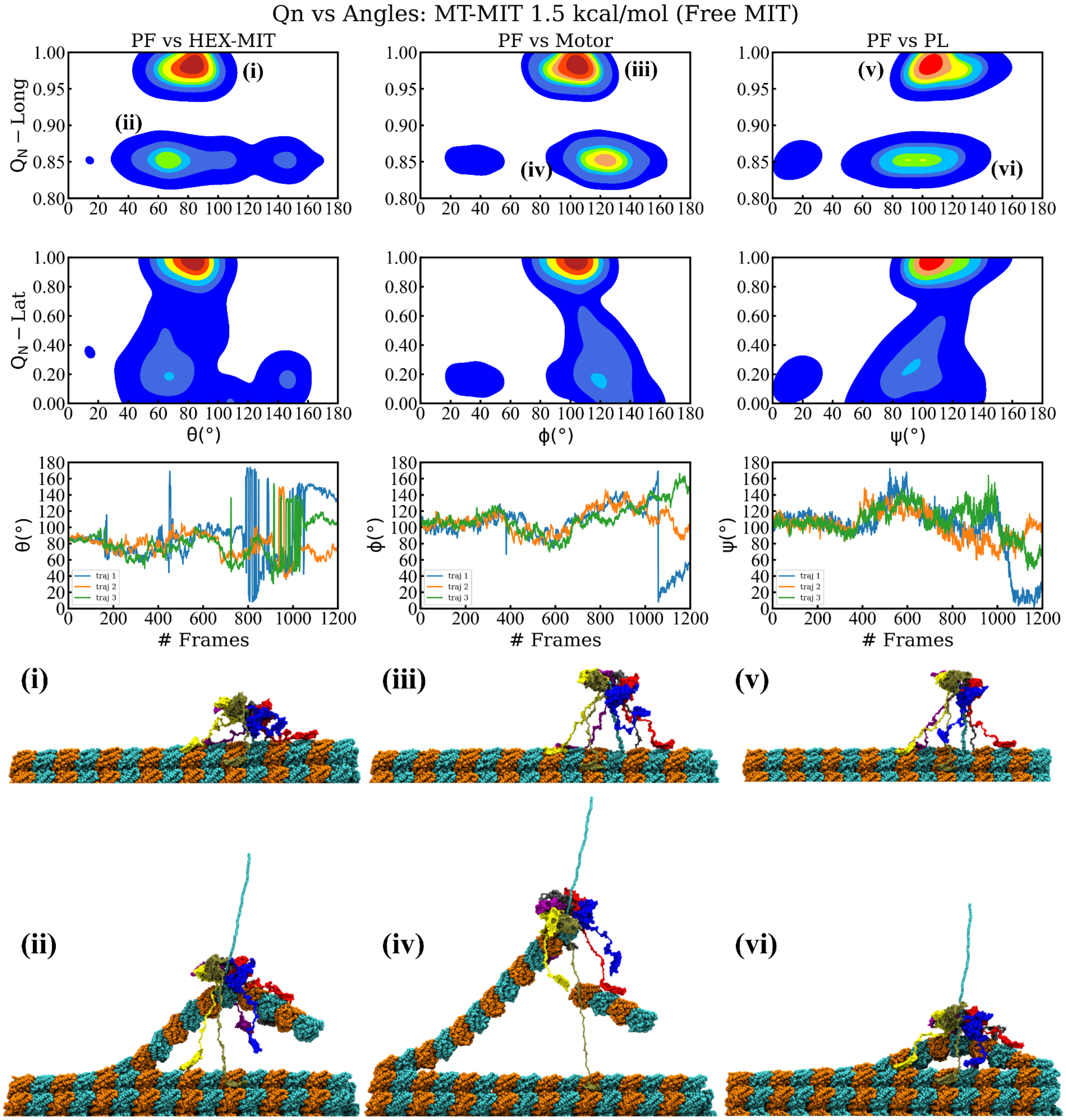

Qn vs Angles: MT-MIT 1.5 kcal/mol (Free MIT)
PF vs HEX-MIT
PF vs Motor
PF vs PL
$Q_N$ − Long
$Q_N$ − Lat
θ(°)
ϕ(°)
ψ(°)
### Frames
traj 1
traj 2
traj 3
(i)
(ii)
(iii)
(iv)
(v)
(vi)

Figure S7: Results of the spastin machine acting on a MT filament for the interaction strength between its MIT domains and the MT lattice set to 1.5 kcal/mol. This is similar to Fig. 2 in the main text.

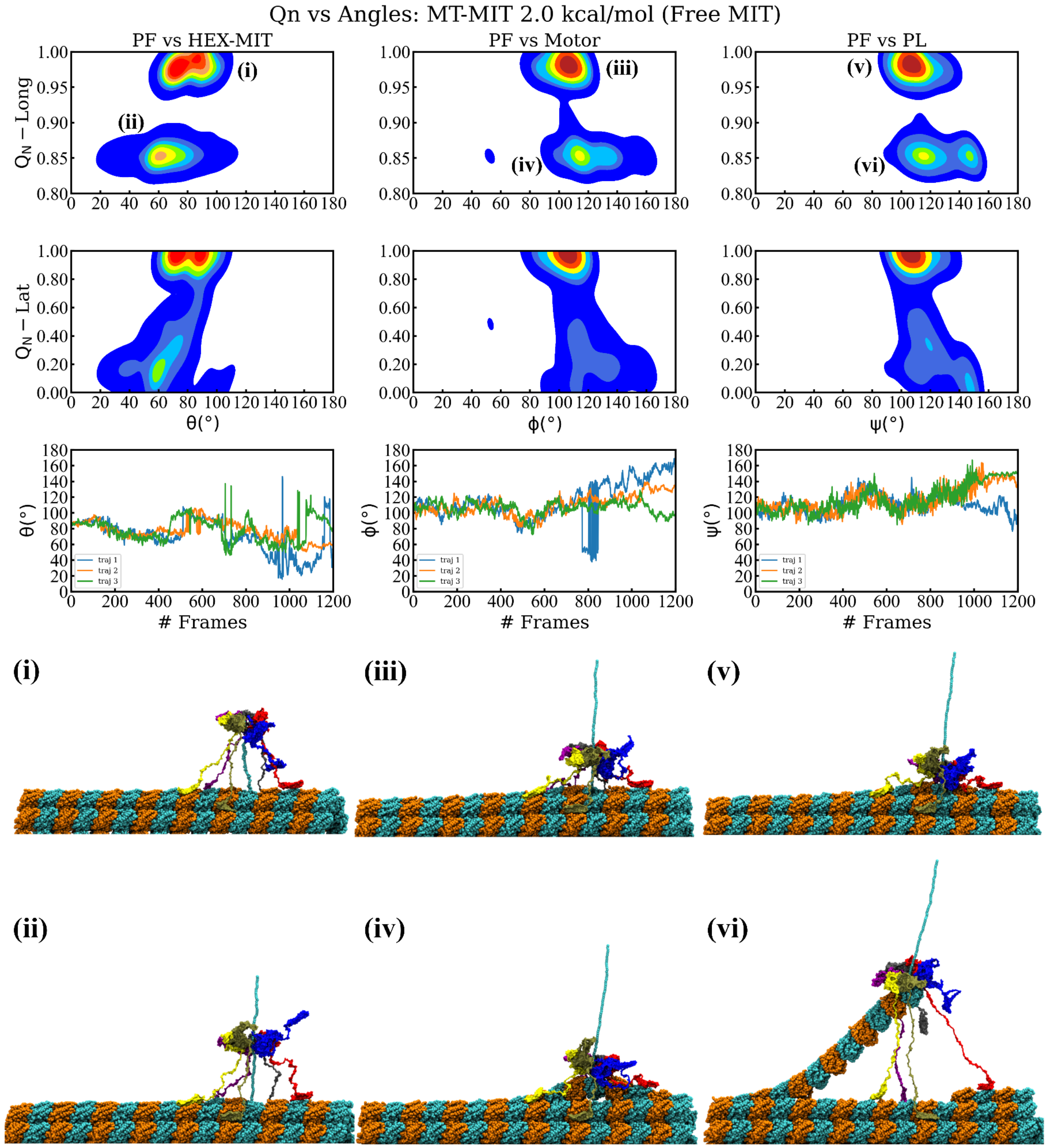
Qn vs Angles: MT-MIT 2.0 kcal/mol (Free MIT)
PF vs HEX-MIT
PF vs Motor
PF vs PL
Q_N − Long
Q_N − Lat
θ(°)
ϕ(°)
ψ(°)
### Frames
traj 1
traj 2
traj 3
(i)
(ii)
(iii)
(iv)
(v)
(vi)

Figure S8: Similar to Fig. S7, for the interaction strength between its MIT domains and the MT lattice set to 2.0 kcal/mol.

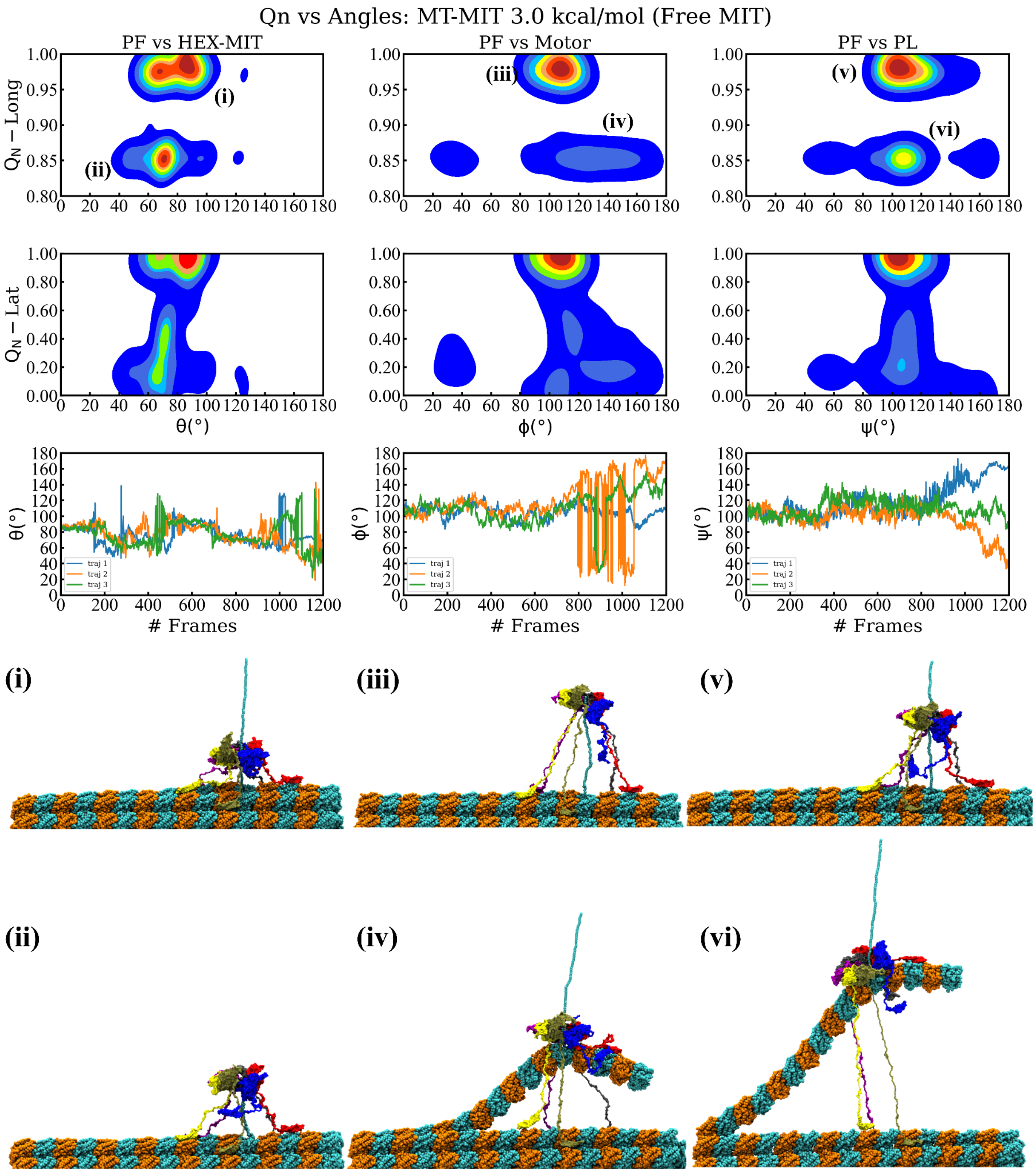

Qn vs Angles: MT-MIT 3.0 kcal/mol (Free MIT)
PF vs HEX-MIT
PF vs Motor
PF vs PL
$Q_N$ − Long
$Q_N$ − Lat
θ(°)
ϕ(°)
ψ(°)
### Frames
traj 1
traj 2
traj 3
(i)
(ii)
(iii)
(iv)
(v)
(vi)

Figure S9: Similar to Fig. S7, for the interaction strength between its MIT domains and the MT lattice set to 3.0 kcal/mol.

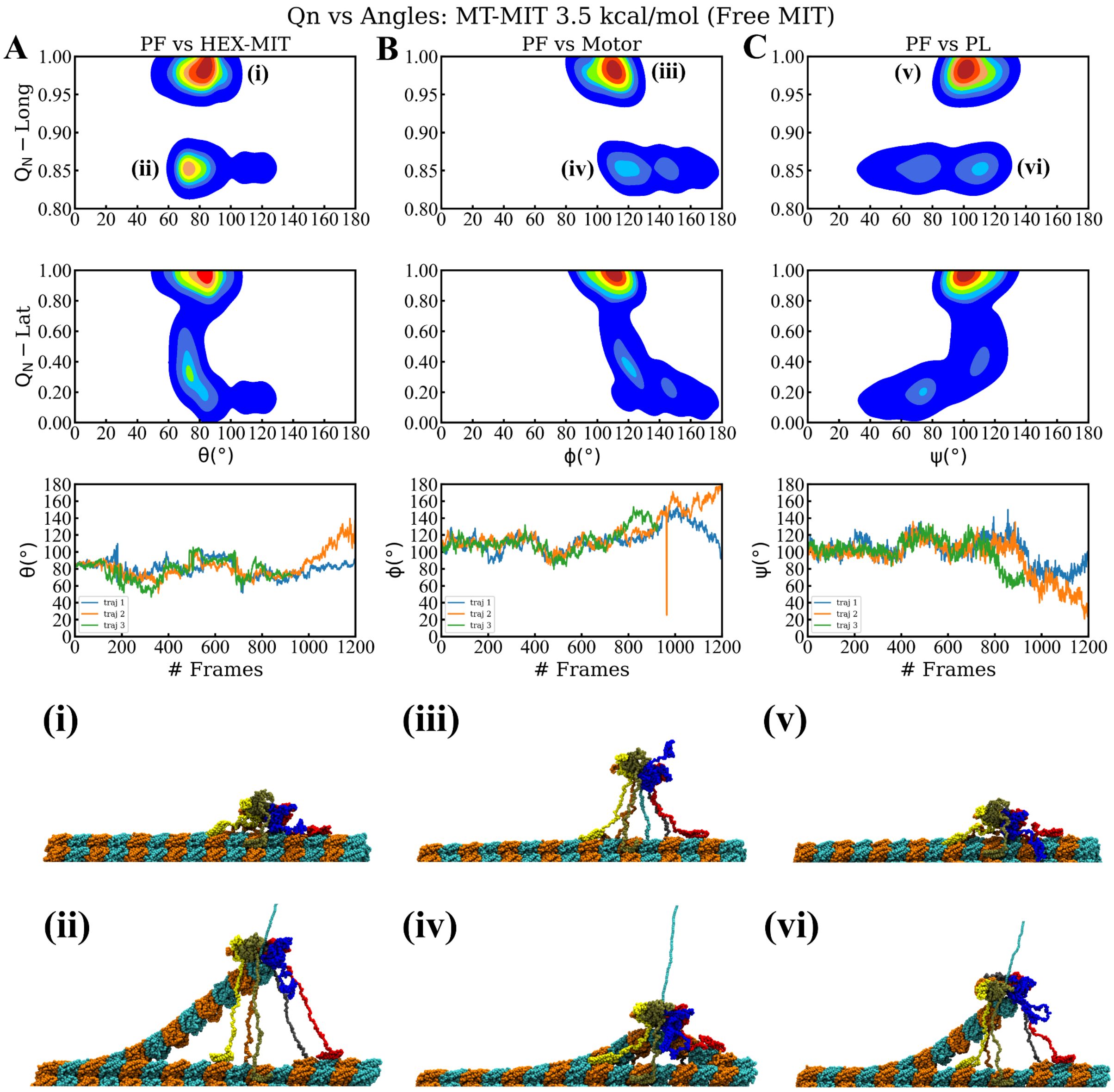

Qn vs Angles: MT-MIT 3.5 kcal/mol (Free MIT)
A
PF vs HEX-MIT
B
PF vs Motor
C
PF vs PL
(i)
(ii)
(iii)
(iv)
(v)
(vi)
$Q_N$ – Long
$Q_N$ – Lat
θ(°)
ϕ(°)
ψ(°)
### Frames
traj 1
traj 2
traj 3

Figure S10: Similar to Fig. S7, for the interaction strength between its MIT domains and the MT lattice set to 3.5 kcal/mol.

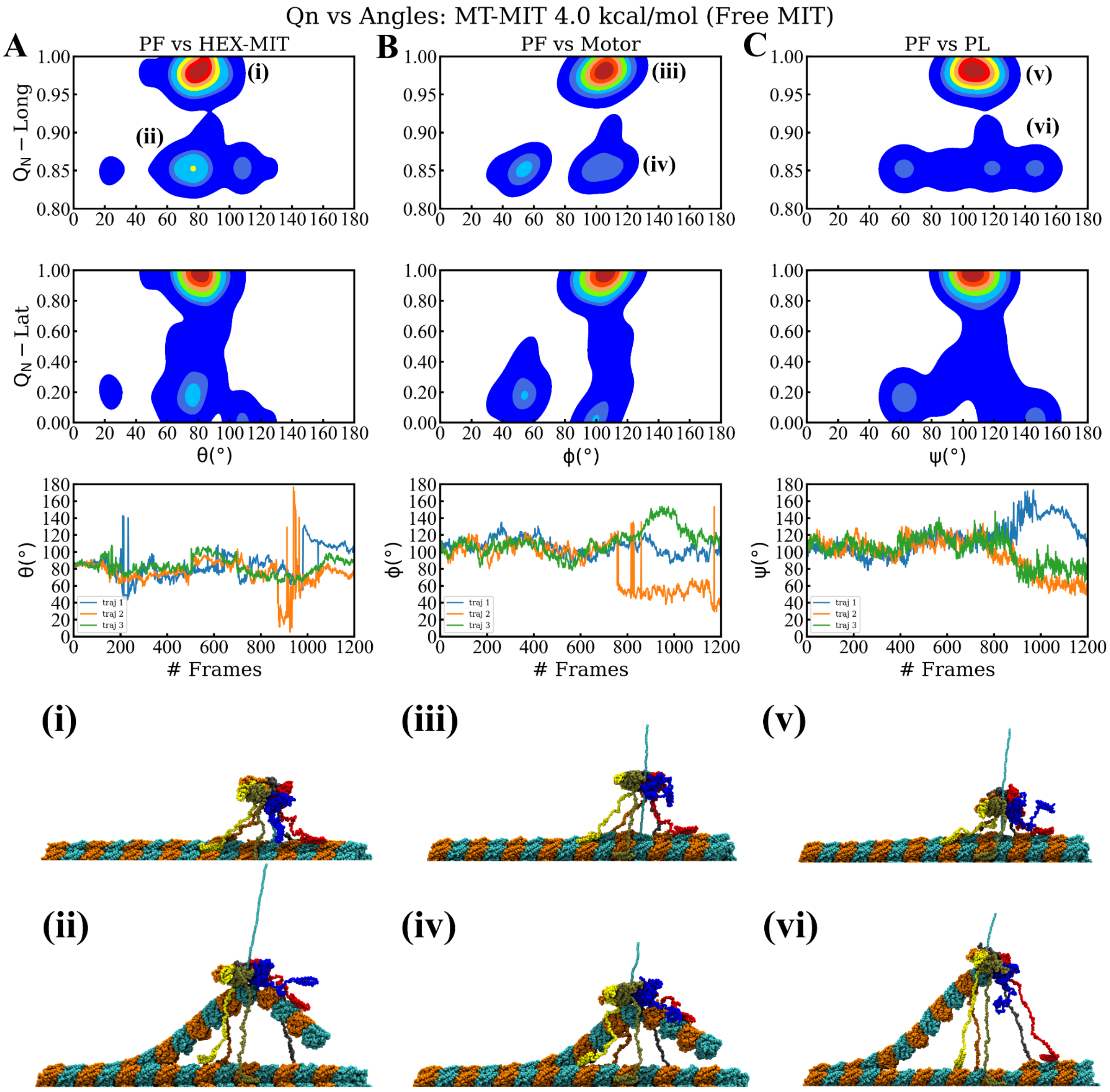
Qn vs Angles: MT-MIT 4.0 kcal/mol (Free MIT)
A
PF vs HEX-MIT
B
PF vs Motor
C
PF vs PL
(i)
(ii)
(iii)
(iv)
(v)
(vi)
$Q_N$ – Long
$Q_N$ – Lat
θ(°)
ϕ(°)
ψ(°)
### Frames
traj 1
traj 2
traj 3

Figure S11: Similar to Fig. S7, for the interaction strength between its MIT domains and the MT lattice set to 4.0 kcal/mol.

**Clustering of DHFR conformations and orientations in ClpY–mediated unfolding and translocation pathways**

Agglomerative hierarchical clustering with complete linkage (top panel) was used to identify the principal conformations and orientations of WT-DHFR and its CP variants along the ClpY-mediated unfolding and translocation pathways. We used the Euclidean metric to evaluate datasets combining the fraction of native contacts $Q_N$ and the polar angle $\theta$. We computed the appropriate cluster numbers,  5 for each setup, using the silhouette[1], Caliski–Harabasz[2], Davies–Bouldin[3] scores.

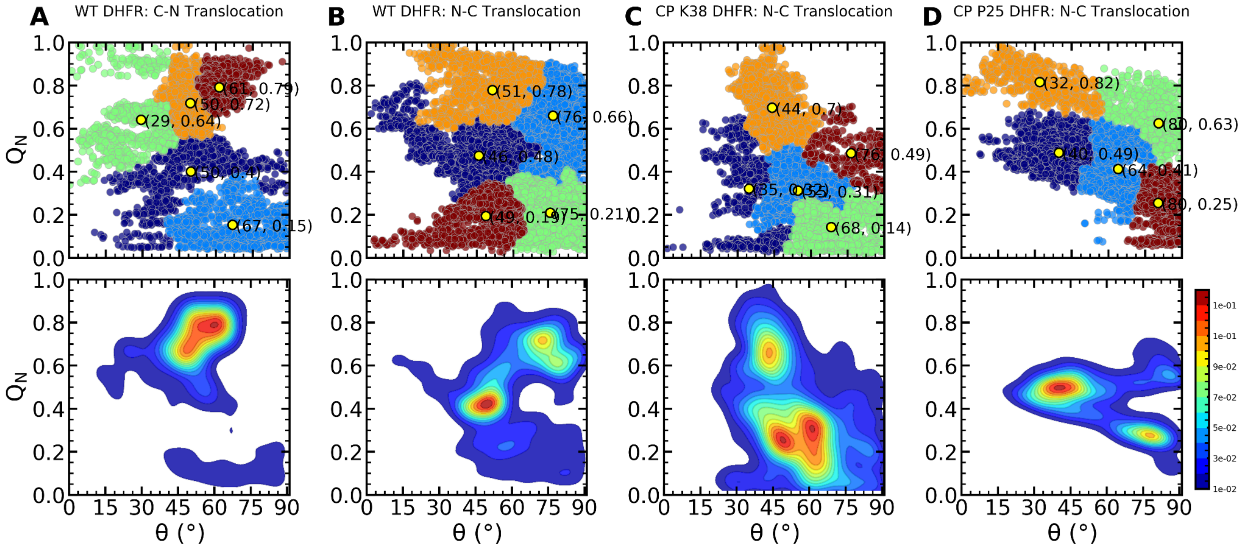


Figure S12: Clustering analysis of DHFR conformations and orientations in ClpY-mediated unfolding and translocation pathways. Principal clusters and probability density maps of datasets of $Q_N$ and the polar angle θ in pathways corresponding to the (A)-(B) WT-DHFR in (A) C-N aand (B) N-C translocation, (C) CP K38 and (D) CP P25. Centroids of clusters are indicated using yellow dots.

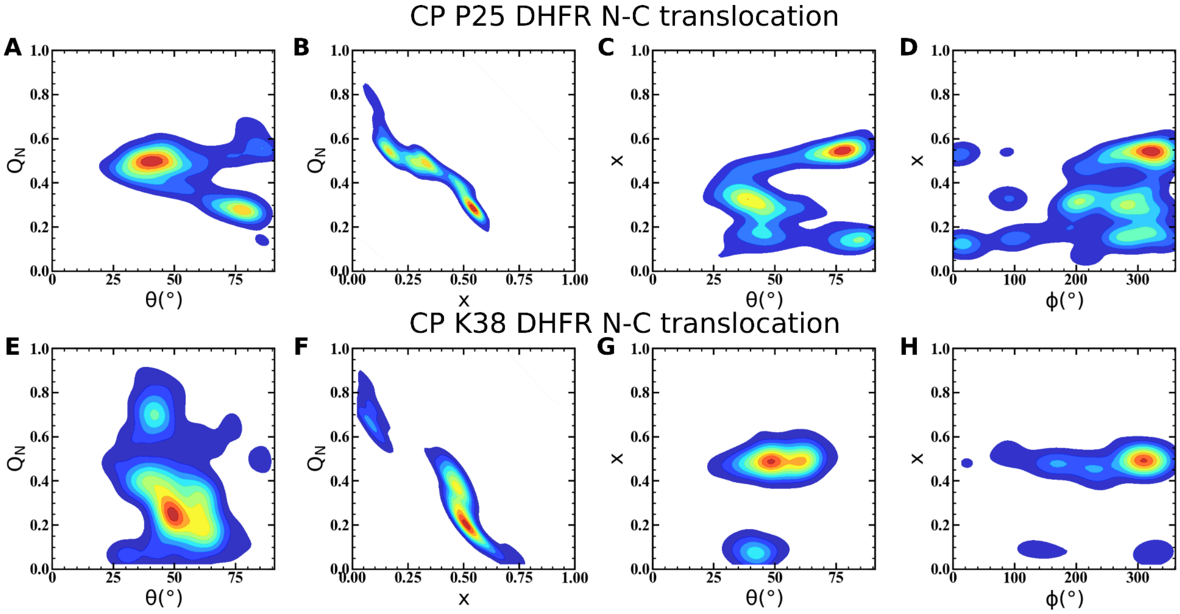


Figure S13: CP DHFR variant orientation at the ClpY pore lumen in unfolding and translocation pathways. Probability density maps of the (A) fraction of native contacts $Q_N$ vs. polar angle θ and (B) translocation fraction x; (C) translocation fraction vs. polar θ and azimuthal φ angles in CP P25 translocation. (E)-(H) Same as in (A)-(D) in the CP K38.
